# Architecture of the Human Default Mode Network: cytoarchitecture, wiring and signal flow

**DOI:** 10.1101/2021.11.22.469533

**Authors:** Casey Paquola, Margaret Garber, Stefan Frässle, Jessica Royer, Yigu Zhou, Shahin Tavakol, Raul Rodriguez-Cruces, Donna Gift Cabalo, Sofie Valk, Simon Eickhoff, Daniel S. Margulies, Alan Evans, Katrin Amunts, Elizabeth Jefferies, Jonathan Smallwood, Boris C. Bernhardt

**Affiliations:** McConnell Brain Imaging Centre, Montreal Neurological Institute, McGill University, Montréal, Quebec, Canada; Institute for Neuroscience and Medicine, INM-7, Forschungszentrum Jülich, Jülich, Germany; Translational Neuromodeling Unit (TNU), University of Zurich & ETH Zurich, Zurich, Switzerland; Max Planck Institute for Cognitive and Brain Sciences, Leipzig, Germany; Institute for Systems Neuroscience, Heinrich Heine Universistät Dusseldorf, Dusseldorf, Germany; Integrative Neuroscience & Cognition Center (INCC – UMR 8002), University of Paris, Centre national de la recherche scientifique (CNRS); Institute for Neuroscience and Medicine, INM-1, Forschungszentrum Jülich, Jülich, Germany; Department of Psychology, University of York, York, United Kingdom; Department of Psychology, Queen’s University, Kingston, Ontario, Canada

## Abstract

The default mode network (DMN) is implicated in many aspects of complex thought and behavior. Here, we leverage *post-mortem* histology and *in vivo* neuroimaging to characterise the anatomy of the DMN to better understand its role in information processing and cortical communication. Our results show that the DMN is cytoarchitecturally heterogenous, containing cytoarchitectural types that are variably specialised for unimodal, heteromodal, and memory-related processing. Studying diffusion-based structural connectivity in combination with cytoarchitecture, we found the DMN contains regions receptive to input from sensory cortex and a core that is relatively insulated from environmental input. Finally, analysis of signal flow with effective connectivity models showed that the DMN is unique amongst cortical networks in balancing its output across the levels of sensory hierarchies. Together, our study establishes an anatomical foundation from which mechanistic accounts of the broad role the DMN plays in human brain function and cognition can be developed.

---

The default mode network (DMN) is a distributed set of brain regions in the frontal, temporal, and parietal lobes with strongly correlated fluctuations^1–3^. It is among the most influential, yet challenging discoveries of modern neuroscience^4,5^. Theories on the role of the DMN initially focused on internally-oriented cognition and its antagonism with “task-positive” networks^6,7^, but increasing evidence shows DMN activity is related to the content of external stimuli^8,9^ and externally-oriented task demands^10–12^. Additionally, DMN subregions can co-fluctuate with regions of “task-positive” networks^13–15^. Thus, the DMN poses a conceptual challenge; how can a neural system be involved in so many different states, particularly since many are seemingly antagonistic, such as perceptually-driven decision making^16^ and perceptually-decoupled cognition^17–19^.

Recent perspectives have argued that resolving the role of the DMN in cognition depends on understanding its anatomy^7,20,21^ because neuroanatomical insights can narrow the search space for conceivable theoretical accounts of its function. While the DMN is typically defined on functional grounds (*i.e.,* strong resting state functional connectivity and relatively lower activity during externally-oriented tasks), its subregions are also connected by long-range tracts^22–24^ and each subregion is maximally distant from primary sensory and motor areas^25^. This topography may allow neural activity in the DMN to be decoupled from perception of the here and now^20^, as neural signals are incrementally transformed across cortical areas from those capturing details of sensory input towards more abstract features of the environment^26,27^. These observations suggest neural activity in the DMN has the potential to be both distinct from sensory input, while also incorporating abstract representations of the external world. This could explain the network’s involvement across diverse contexts^20^. Although this topographical perspective, in principle, accounts for its broad involvement in human cognition, we lack a detailed explanation of how the neural circuitry within the DMN enables this hypothesised role^28^.

Given the highly distributed nature of the nodes of the DMN, it is likely to be heterogeneous in terms of its microarchitecture, however, the specific nature of this heterogeneity remains unknown. This gap in our knowledge prevents us from leveraging anatomical insights to better understand the DMNs connectivity and function. On the one hand, it is conceivable that regional differences in the DMN are most pronounced between subregions situated in different lobes, with different white matter tracts connecting each subregion^29,30^. On the other hand, an increasing literature has emphasised the presence of large-scale cytoarchitectural gradients across the cortex, suggesting a microstructural differentiation between sensory and transmodal regions as well as long distance similarities in microarchitectural profiles^31–33^. Such large-scale cytoarchitectural gradients can also underlie organisation within a subregion, such as the mesiotemporal lobe and insula^34,35^. Thereby, fine-grained intra-regional differentiation is another important contributor to heterogeneity within the DMN. Fine-grained patterns of differentiation need not be gradients, however. Primate tract-tracing and precision functional imaging studies have revealed interdigitation of connectivity within regions of the DMN, such as the prefrontal cortex and the inferior parietal lobe^36–38^. Thus, while laminar connectivity across the cortex follows consistent rules^39,40^, microstructure and connections can be organised locally in a range of patterns from relatively smooth gradients to chequered interdigitation. Recent innovations in whole-brain human histology as well as quantitative *in vivo* MRI at high fields have made it possible to determine how these various findings manifest within the DMN, enabling the derivation of an anatomically grounded blueprint of its organisation.

The microarchitectural make-up of the DMN ultimately influences how it processes information because microarchitecture influences both the intrinsic computation within a region and its connectivity to other regions; the two sides of functional specialisation. For instance, the degree of laminar differentiation, which varies in a graded manner across the cortex^31,41^, reflects different specialisations of the underlying cortical microcircuits, ranging from externally-focused sensory areas through unimodal and heteromodal cortex to amodal agranular areas^42,43^. Patterns of projections also systematically vary along this gradient^40,44–46^, forming a hierarchical architecture of cortico-cortical tracts spanning from primary sensory areas to the prefrontal cortex and mesiotemporal lobe^39,47,48^. Whether a hierarchy constrains connectivity within association cortex (such as the DMN) has been questioned, however^2,49,50^. Instead, the DMN may comprise densely interconnected yet spatially distributed circuits, operating in parallel to the canonical sensory hierarchies^49^. Distinguishing between hierarchical and non-hierarchical schemas relies upon characterising how signal flows with respect to the underlying microarchitecture. To this end, state-of-the-art connectivity mapping approaches that emphasise directed signal flow, including recently introduced measures of navigation efficiency of structural connections ^51^ and regression dynamic causal modelling of functional signals^52,53^, can help adjudicate between different theoretical perspectives. In combination with data-driven microarchitectural mapping, these approaches can elucidate how cortical anatomy constrains the communication of the DMN, shedding light on the perhaps unique organisational principles of human association cortex.

Here, we capitalise on a combination of *post-mortem* histology and multimodal *in vivo* neuroimaging to map DMN microarchitecture and examine how microarchitecture contributes to its structural and functional embedding in the brain. In particular, we leverage (i) an established atlas of cytoarchitectural taxonomy (“cortical types”)^31,54^, (ii) whole-brain 3D histology for fine-grained cytoarchitectonic mapping^55,56^ and (iii) multimodal *in vivo* neuroimaging for approximations of structural wiring and functional flow. Finally, (iv) using high-field 7 Tesla (7T) MRI, we demonstrate how the discovered relationships between microarchitecture, connectivity and function of the DMN exist within an individual.

## Cytoarchitectural heterogeneity

The DMN is generally agreed to encompass subsections of the (1) parahippocampal cortex, (2) precuneus and posterior cingulate cortex, (3) a caudal region of the inferior parietal lobule, (4) middle temporal cortex, (5) inferior fronto-lateral cortex, and (6) a region of the prefrontal cortex, primarily covering the superior frontal gyrus and anterior cingulate, as well as a small part of the middle frontal gyrus^7,57,58^. Throughout our primary analyses, we use the most common atlas of the default mode network^2^ (**Figure 1A**) and identified six spatially contiguous subregions within each hemisphere that correspond to the abovementioned regions (see **Supplementary Table 1** for Von Economo areas and Schaefer parcels encompassed by each subregion). In supplementary analyses, we show the replicability of key findings with alternative delineations of the DMN, specifically based on deactivations during externally-oriented tasks^20^, independent component analysis of task-based fMRI^59^, and individualised Bayesian modelling of functional communities^60^.

**Figure 1:**
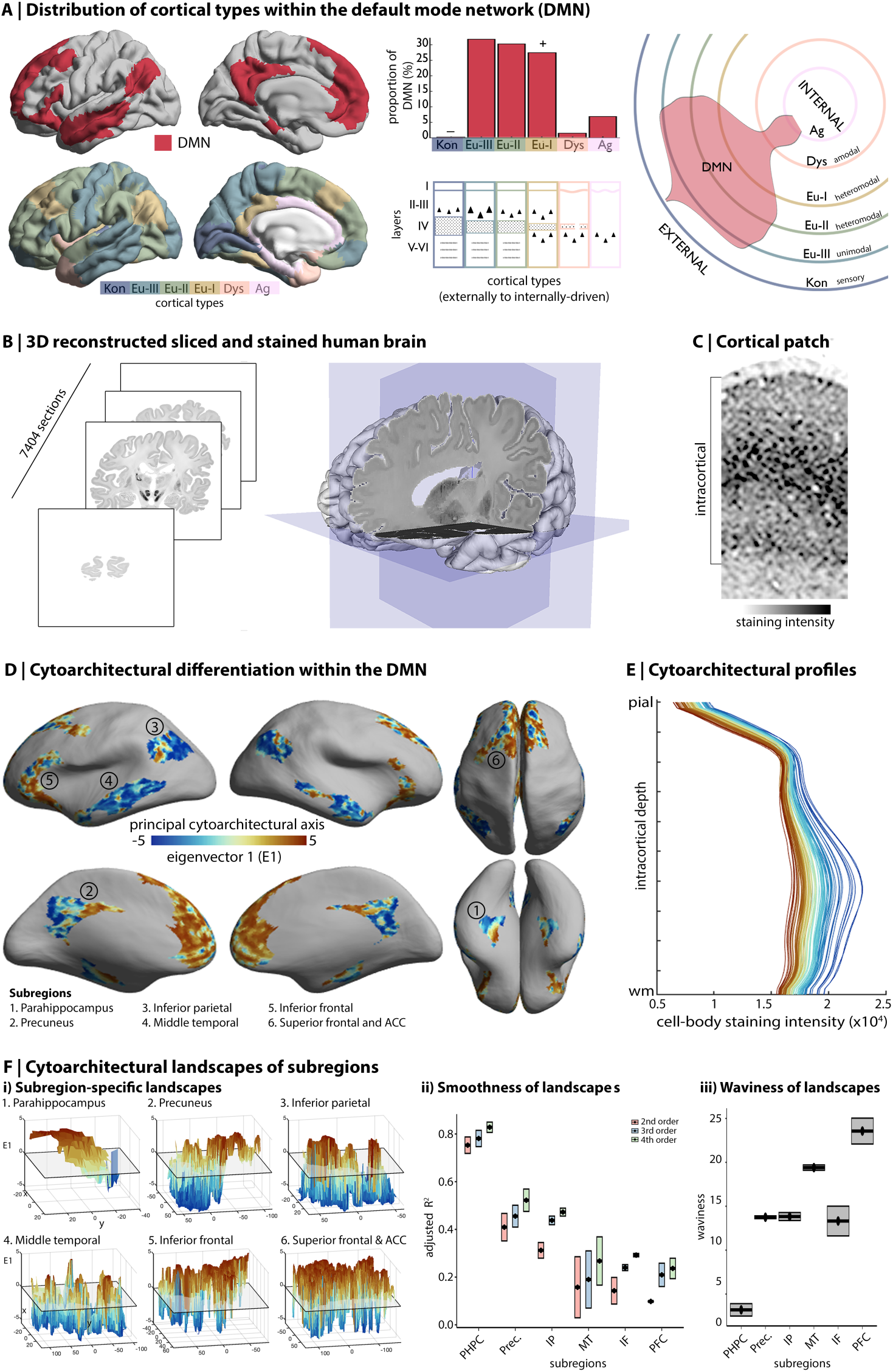
Cytoarchitectural heterogeneity of the DMN. **A)** *Upper left*. The most common atlas of the DMN^2^, used in primary analyses and shown on the cortical surface. *Lower left.* Cytoarchitectonic atlas of cortical types^31,54^. *Upper middle.* Histogram depicts the frequency of cortical types within the DMN. + is indicative of significant over-representation and – is under-representation, relative to whole cortex proportions. *Lower middle.* The schematic highlights prominent features that vary across cortical types, including the location/size of largest pyramidal neurons (triangles), thickness of layer IV, existence of sublayers in V-VI (grey dashed lines), regularity of layer I/II boundary (straightness of line). Kon=koniocortical. Eul=eulaminate. Dys=dysgranular. Ag=agranular. *Right.* Circular plot represents the spread of the DMN from externally-to internally-driven cortical types. The percentage of each type within the DMN is depicted by the amount of the respective line (not the area in between lines) covered by the red shaded violin. See also ^21,43,63^ for similar schematics. **B)** 7404 coronal slices of cell-body-stained sections (20 μm thickness) were reconstructed into a 3D human brain model, BigBrain^55^. **C)** Example cortical patch shows depth-wise variations in cell-body-staining in BigBrain. **D)** The principal eigenvector (*E1*) projected onto the inflated BigBrain surface shows the patterns of cytoarchitectural differentiation within the DMN. **E)** Line plots represent cell-body-staining intensity by intracortical depth (from pial to white matter boundary) at different points along E1. Cortical points with lower E1 (*blue*) have peaked cellular density in mid-deep cortical layers, indicative of pronounced laminar differentiation, whereas cortical points with higher E1 (*red*) have more consistent cellular density across cortical layers, illustrating lower laminar differentiation. **F)** The topography of E1 in each subregion shown as 3D surface plots, with E1 as the z-axis. The x- and y-axes are defined by Isomax flattening of each subregion. Left boxplots show the proportion of variance in E1 explained by spatial axes (x,y) for each subregion and for models of increasing complexity (2^nd^-4^th^ order polynomial regression). Boxplot range depicts hemisphere differences in adjusted R^2^, while the centre point is the adjusted R^2^ averaged across hemispheres. Right boxplots show waviness of E1 in each subregion. Together, these metrics, further described and validated in **Supplementary Figure 5**, quantify how cytoarchitectural landscapes vary between subregions; from a relatively simple gradient in the parahippocampus, well-explained by the spatial regression model and with low waviness, to marked fluctuations in the dorsal prefrontal cortex, characterised by high waviness and poor regression model performance. PHPC=parahippocampus. Prec.=precuneus. IP=inferior parietal. MT=middle temporal. IF=inferior frontal. PFC=prefrontal cortex.

The most noticeable difference in cytoarchitecture across cortical regions is the degree of laminar differentiation *i.e.,* the distinguishability and thickness of layers. Degree of laminar differentiation is highest in primary sensory areas and decreases along the cortical mantle in a graded manner, reaching a low in agranular cortex, which neighbours hippocampal and piriform allocortex. This gradient of laminar differentiation is synopsised by six cortical types, originally defined by Von Economo^31,54^(**Figure 1A)**. Patterns of projections also systematically vary along this gradient^40,44–46^, forming a hierarchical architecture spanning primary sensory areas to the prefrontal cortex and hippocampus^39,47,48^. Notably, the cortical types (synonymous with the levels of sensory hierarchies) are hypothesised to reflect different specialisations of the underlying cortical microcircuits, ranging from externally-focused sensory areas through unimodal and heteromodal cortex to agranular, paralimbic areas^42,43^. This hypothesised relationship, based primarily on neurophysiological evidence in non-human primates and lesion studies in humans^43,61^, is supported here by meta-analytical decoding of the cortical types, using activation maps from thousands of functional MRI studies (**Supplementary Figure 1**).

Based on overlap of the DMN atlas with a cytoarchitectonic atlas of cortical types^31,54^, we found the DMN contains five of six cortical types (**Figure 1A**). This make-up was distinctive, relative to other functional networks (**Supplementary Table 2,** all Kolgomorov-Smirnoff tests>0.11, p<0.001). Indeed, pair-wise comparisons showed that all networks exhibited a unique composition of cortical types (**Supplementary Figure 2**). Notably, of all functional networks, the DMN contains the most balanced representation of the three eulaminate types, which are commonly associated with processing of sensory information and its progressive integration (eulaminate-I, II, and III). In addition, the DMN contains dysgranular and agranular cortex that are often linked to internally-generated processes, such as memory and affect^43,62^ (**Supplementary Figure 1**). These cortical types are not equally represented within the DMN, however (χ^2^=1497, p<0.001). Approximately 90% of the DMN is eulaminate, which is even higher than the cortex-wide rate of 84% (**Supplementary Table 2**). To evaluate whether this type of cortical is over-represented in the DMN, we compared the proportion of cortical types within the DMN and within 10,000 rotated versions of the DMN. The rotated versions are generated by randomly spinning the functional network atlas on a spherical representation of the cortex, providing a null distribution of outcome statistics that account for the network’s size and distribution. In doing so, we found that the DMN over-represents eulaminate-I (18% increase; p_spin_=0.006), classically known as “heteromodal” cortex, which is hypothesised to process information from multiple sensory domains^43^ (**Supplementary Figure 1**). This distinctive composition of cortical types was evident regardless of slight alterations to the DMN atlas (**Supplementary Figure 3**). The broad range of cortical types in the DMN, combined with the over-representation of eulaminate-I, is consistent with a role of this network in integration of information from multiple systems including those linked to sensory and memory processes.

Having established the DMN contains a broad array of cortical types, we next adopted a data-driven approach to characterise fine-grained spatial patterns of cytoarchitectural variation. We transformed the functional network atlas^2^ to a 3D cell-body-stained *post-mortem* human brain^55^ using specially tailored cortical registration procedures^56,64^. Using intracortical profiles of cell-body-staining intensity (**Figure 1C,E**), we assessed cytoarchitectural variability within the DMN, mapping cytoarchitectural variation via unsupervised non-linear manifold learning^65^ (**Figure 1D**, see also **Supplementary Figure 4**). The first eigenvector of this manifold (E1), hereafter referred to as the *cytoarchitectural axis*, described a shift in the shape of the underlying cytoarchitectural profiles from peaked to flat (**Figure 1E**) and reflects differences in how cellular density varies within the cortex (**Figure 1C**). The cytoarchitectural axis is anchored on one end by unimodal eulaminate-III cortex (*e.g.,* retrosplenial and posterior middle temporal) and on the other by agranular cortex (*e.g.,* medial parahippocampus and anterior cingulate). Thus, the endpoints of the cytoarchitectural axis are the most extreme cortical types found within the DMN (**Supplementary Figure 4**). Beyond the endpoints, however, the cytoarchitectural axis deviates from the gradient described by cortical types^31,33,43^ (**Supplementary Figure 4**). This pattern does not discriminate subregions of the DMN or follow an anterior-posterior gradient as seen in neuronal density^66^. Instead, we observed a mosaic of different spatial topographies across DMN subregions, where neighbouring points are sometimes distinct and distant points are sometimes similar. Our data-driven approach thus indicates that organisation within the DMN is unlike those observed across sensory hierarchies and is relatively unconstrained by large-scale spatial gradients^49,50^.

A closer look at the topography of cytoarchitecture highlights the (dis)similarity of neighbouring areas within the DMN. Given the ubiquity of local connectivity in the cortex^67,68^, topography provides important information on the form of communication within spatially-contiguous subregions. Subregions of the DMN evidently vary in terms of their cytoarchitectural topography (**Figure 1F**), and we quantified these differences using two complementary measures: smoothness and waviness. The smoothness of the microarchitectural landscape was calculated by evaluating the proportion of variance in the cytoarchitectural axis that could be accounted for by spatial axes. Waviness was indexed by deviations from the mean, a common technique in mechanical engineering^69^ (see **Supplementary Figure 5** for simulation-based validation of these metrics). We found that subregions significantly differ in terms of both smoothness and waviness (smoothness: 2^nd^/3^rd^/4^th^ order; F=14.5/14.9/20.1, p<0.004; waviness: F=48.3, p=0.001). Smoothness was particularly high in the parahippocampus, showing that its cytoarchitectural axis follows a relatively smooth gradient, as may be predicted from previous anatomical research^35,70^. Conversely, the prefrontal cortex exhibits especially high waviness. This pattern of frequent changes across the cortex, back-and-forth between two contrasting properties, is notably reminiscent of “interdigitated” connectivity patterns that are known to exist within the prefrontal cortex^36,37,71^. This analysis establishes that the DMN contains distinct cytoarchitectural patterns representative of two different ways that neural signals are hypothesised to be integrated in the cortex: A mesiotemporal gradient associated with progressive convergence of information^72,73^, and prefrontal interdigitation that enables information from disparate sources to be linked^36^.

## Receivers on the periphery and an insulated core

Thus far, our analyses provided evidence that the DMN contains highly variable types and arrangements of cortical microarchitecture, findings consistent with the hypothesis that a wide range of neural signals can be integrated within the regions that make up this network. Next, we explored how the anatomical features of the DMN relate to its connectivity using multi-modal MRI. We hypothesised that connectivity would co-vary with the cytoarchitectural axis (E1, **Figure 1D**), because propensity for connectivity increases with cytoarchitectural similarity^44,74,75^. While this principle has been observed across association and sensory regions^38,45,71^, it remains unclear whether, and how, it applies specifically to the connectivity of the regions within the DMN.

First, we measured communication efficiency along white matter tracts^51^ using diffusion magnetic resonance imaging (MRI) tractography^76^. Navigation is a decentralised communication strategy that is particularly suited to spatially embedded networks, which has recently been proposed to study structural connectivity and structure-function relationships in the human brain ^76^. Navigation involves identifying a single, efficient path between two nodes, without assuming global knowledge of network topology, thus overcoming a key shortcoming of other communication models, such as the shortest path. Instead, navigation uses physical distances between nodes to determine the path. Specifically, starting from a source node, the path is extended to a neighbour (*i.e.,* a structurally connected node) that is closest in physical space to the final target node. In doing so, the approach can achieve a balance of local clustering and long-range connectivity that is characteristic of widely distributed brain networks, such as the DMN. We found that the propensity to communicate with other cortical areas (indexed by average “navigation efficiency”^51^, see *Methods* for details) varied within the DMN [coefficient of variation (CoV)=18%]. Areas towards one end of the DMN’s cytoarchitectural axis, specifically those with more peaked cytoarchitectural profiles, such as the anterior cingulate and more anterior aspect of the precuneus, exhibited more efficient communication with the rest of the cortex (r=-0.60, p_spin_=0.001, **Figure 2Ai**). This effect was particularly pronounced for communication with perceptually-coupled cortical types (koniocortical/eulaminate-III/eulaminate-II; r=-0.63/-0.60/-0.38, p_spin_<0.025, **Figure 2Ai**). Thus, the organisation of the DMN, revealed by cytoarchitectural analysis, also correlates with spatial patterns of how tract-based communication varies within the DMN, especially between the DMN and cortical areas engaged in sensory processing. This pattern of covariation was specific to connectivity between the DMN and non-DMN areas, and did not apply to connectivity within the DMN (**Supplementary Figure 6**), suggesting that inter- and intra-network connectivity may involve distinct rules of organisation that are embedded within in more general, cortex-wide principles, such as the structural model^40,77^.

**Figure 2:**
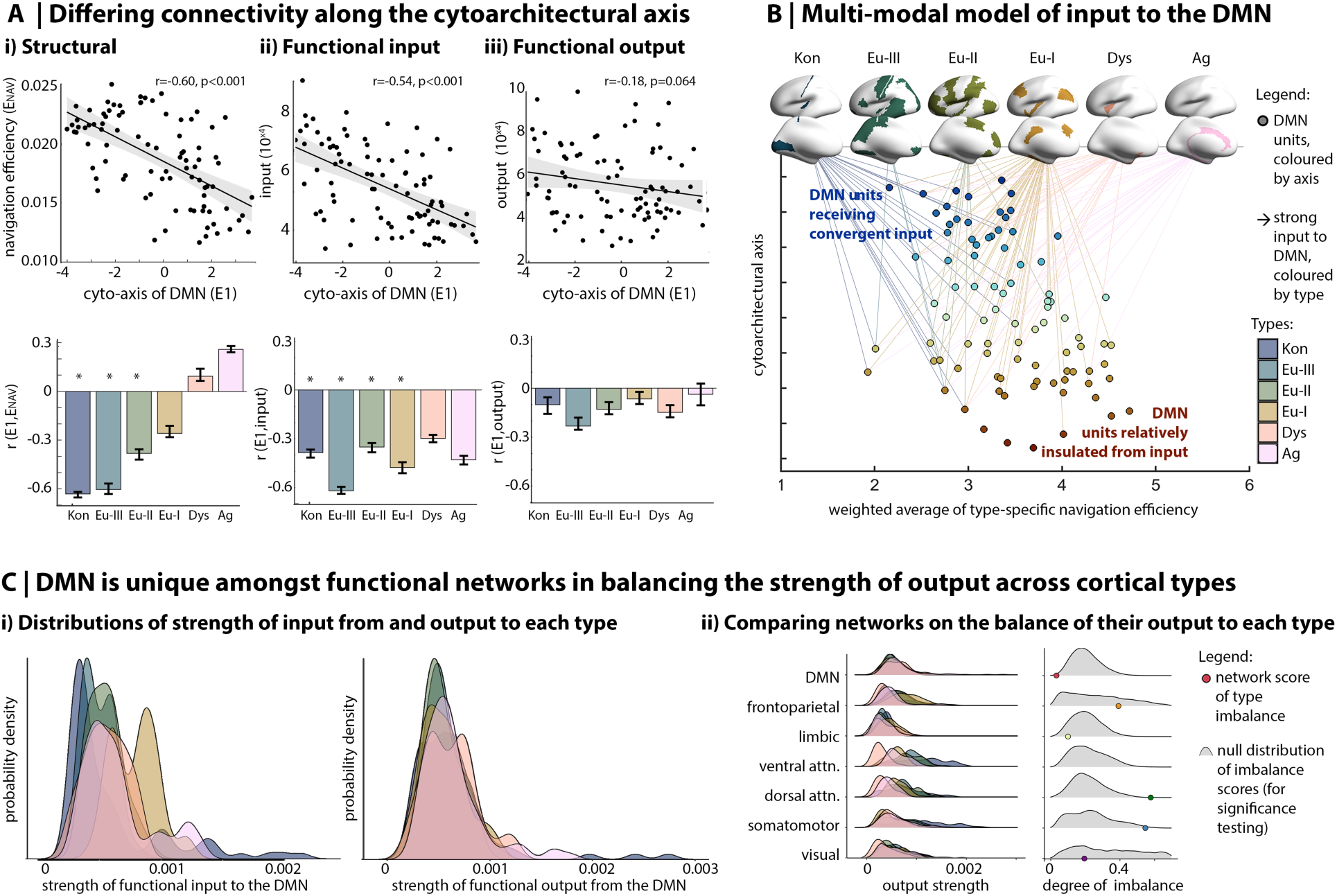
Organisation of DMN connectivity. **A)** *Above.* Scatterplots show the correlation of the cytoarchitectural axis (E1) with average (i) structurally-modelled navigation efficiency, (ii) functionally-modelled input and (iii) functionally-modelled output. Each point represents a node of the DMN. Their subregion assignment is illustrated in **Supplementary Figure 6A**, **Supplementary Figure 7** and **Supplementary Table 1**. *Below.* Bar plots shows the linear correlation coefficient (r)of E1 with average connectivity to each cortical type. The stability of the correlation coefficient was calculated by repeating the procedure in 10 folds, each including 90% of datapoints. Error bars indicate the standard deviation of the r value across folds. Significant (*) negative r values indicate that DMN nodes with peaked profiles have (i) higher navigation efficiency with externally-driven cortical types, and (ii) stronger input from most cortical types. Kon=koniocortical. Eul=eulaminate. Dys=dysgranular. Ag=agranular. **B)** Multi-modal model of DMN organisation shows the dual character of the DMN, including areas with convergent input and insulated areas. All points in the scatterplot represent units of the DMN, are coloured by position along the cytoarchitectural axis (also y-axis) and are organised along the x-axis based on weighted average of type-specific navigation efficiency. Top 75% of functionally-defined inputs are shown. **C)** (i) Coloured ridge plots show probability distributions of connectivity between the DMN and each cortical type. Notably, for functional output the DMN exhibits overlapping, normal distributions, whereas for functional input, type-wise differences are evident. **ii.** Focusing on functional output, coloured ridge plots show distributions for all networks, illustrating more balance between types in the DMN. *Right.* The imbalance of connectivity to distinct cortical types was evaluated as the Kullback-Leibler (KL) divergence from a null model with equal connectivity to each type. The coloured dots show the empirical KL divergence for each network and the grey density plots show the null distribution of KL divergence values based on 10,000 spin permutations. Permutation testing indicated that the DMN is unique among functional networks in balancing output across cortical types (*i.e.,* imbalance lower than 95% of permutations).

Next, we examined the consequences of this structural organisation on the functional flow of information in the cortex. We applied regression dynamic causal modelling, a scalable generative model of effective connectivity^52^ to resting state fMRI timeseries of 400 isocortical parcels^78^, covering the entire isocortex. Effective connectivity aims to describe directed interactions among brain regions, with estimates describing how different regions influence each others’ timeseries. Typically, effective connectivity parameters are estimated in a Bayesian framework by solving a set of differential equations in the time domain (*i.e.,* classic DCM), but computational cost of model inversion limits the number of regions that can be included. rDCM overcomes this limitation by converting the equations into an efficiently solvable Bayesian linear regression in the frequency domain. In doing so, rDCM allows computation of effective connectivity parameters for hundreds of brain regions. In prior work, the face and construct validity of rDCM for inferring effective connectivity parameters during resting state has been established using comprehensive simulations and by comparing rDCM against alternative generative models of rs-fMRI data for small networks^79^. In the current work, we conducted a whole-cortex rDCM, then selected DMN parcels as targets for “functional input” analyses and DMN parcels as seeds for “functional output” analyses. Functionally estimated input and output varied within the DMN (CoV=24% and 29%, respectively). Average strength of input was significantly higher to those areas of the DMN with more peaked cytoarchitectural profiles (r=-0.54, p_spin_<0.001), *i.e.,* those regions that were also highlighted as having more efficient communication with the rest of the cortex in the above structural connectivity analysis (see **Supplementary Figure 7** for a comparison of cortical maps).

Examination of type-specific connectivity showed limited discrimination between cortical types, whereby inputs from externally- and internally-focused cortical types were all concentrated on DMN areas with peaked cytoarchitectural profiles (**Figure 2Aii-iii, Supplementary Table 3**). Thus, multiple inputs converge upon a subset of DMN subunits, such as inferior parietal and precuneus areas, while a subset of DMN subunits, those with flat cytoarchitectural profiles, remained relatively insulated from cortical input. Output did not co-vary with the cytoarchitectural axis (r=-0.18, p_spin_=0.064, **Figure 2Aii-iii**). These findings were consistent in a replication dataset and when including subcortical structures and the hippocampus in the model (**Supplementary Table 3**). Together, these analyses suggest that the DMN comprises two microarchitecturally distinct subsets – one with highly efficient tract-based communication with cortical areas implicated in perception and receiving convergent input from across all levels of sensory hierarchies, while another that exhibits less efficient tract-based communication with the rest of the cortex and is relatively insulated from input signals from sensory systems (**Figure 2B**).

## A UNIQUE BALANCE OF OUTPUT

Focusing on the anatomy of the DMN revealed its distinctive pattern of cytoarchitectural heterogeneity, which constrains how it communicates with other systems. Now, we turn our attention to how these anatomical properties contribute to the position of the DMN in the large-scale functional organisation of the cortex by understanding how effective functional connectivity of the DMN is distributed across cortical types.

First, we discovered that the DMN communicates in a balanced manner with all cortical types. Compared to all other functional networks, the DMN exhibits the most balanced efficiency of communication across cortical types (*i.e.,* lowest KL divergence from null model, **Supplementary Figure 8**, see **Supplementary Table 4** for full statistics). We could further specify that output of the DMN is balanced across the cortical types, but input is not (**Figure 2Ci**, see **Supplementary Table 4** for full statistics and replication). In other words, the DMN outputs signals in approximately equal strength to all cortical types (*i.e.,* all levels of sensory hierarchies). Of all the functional systems in the human cortex, only the DMN exhibited this balance in output across cortical types (**Figure 2Cii**). The spatial distribution, internal heterogeneity, and connectivity of the DMN, thus, engender a unique ability to receive temporally distinct signals and then send neural signals that influence all levels of the sensory hierarchies in a similar manner.

## Correspondence of microarchitecture and connectivity within an individual

To demonstrate that our findings generalise to single individuals, we acquired high-resolution quantitative T1 (qT1) relaxometry MRI, alongside diffusion weighted and functional MRI in eight healthy individuals using a 7 Tesla MRI system. Methods were identical to those described above, except that histology was replaced by qT1. We hypothesised that qT1, sensitive to cortical myelin^80,81^, could recapitulate regional differences in cytoarchitecture, because cortical areas and intracortical layers defined on cyto- or myelo-architecture align^82,83^, and our previous work has shown strong correspondence of principal axes of microstructural differentiation derived from histology and qT1 MRI^33^. While the qT1 and histological datasets differ in terms of biological sensitivity (myelin *vs* cell bodies) and resolution (500μm *vs* 100μm), the patterns of microarchitectural differentiation in the DMN had moderate similarity between the modalities (r_avg_=0.34, p_avg_<0.001), for example highlighting microstructural differences of the prefrontal cortex from the lateral temporal region (**Figure 3A**). We also repeated the analysis using individual-specific DMNs (see Methods, ^84^) and found highly similar axes (**Supplementary Figure 9**). Thereby, microstructural variation within the DMN is not due to idiosyncratic positioning of the DMN, relative to the group-average atlas.

**Figure 3:**
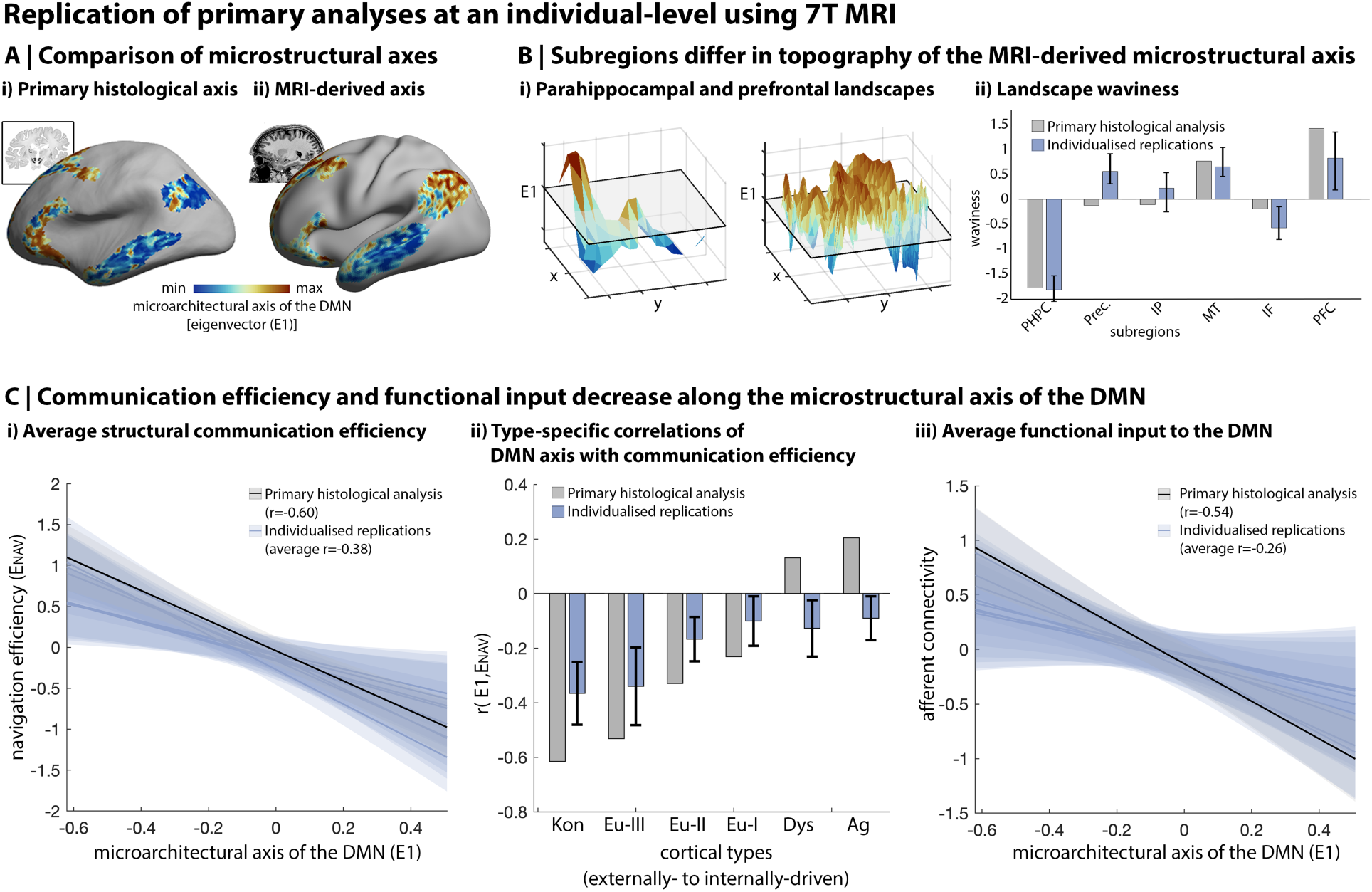
**A)** The principal eigenvector of microstructural variation in the DMN (E1) was extracted from myelin-sensitive quantitative MRI (qT1)^80^, in line with the procedure employed on the histological dataset (“BigBrain”), revealing similar patterns. **B)** The roughness of MRI-derived microstructural differentiation varied between subregions in line with histological evidence. The parahippocampus exhibited a graded transition from high-to-low E1, reflected by high smoothness and low waviness, whereas the prefrontal cortex exhibited an undulating landscape with high waviness. **C)** Using individual-specific measures, we consistently found that cortical points with higher E1 were associated with *(left)* lower average navigation efficiency, (*centre*) especially lower navigation efficiency with perceptually-coupled cortical types, and (*right*) lower functional input. Thus, in line with histological evidence, the MRI-based approach highlights that a subsection of the DMN is relatively insulated from external input. Line plots are presented with 95% confidence interval shading.

While idiosyncrasies and cross-modal differences were evident, especially in the lateral parietal and anterior cingulate regions (**Supplementary Figure 9**), the topography of microarchitectural differentiation was similar in both qT1 and histological datasets, varying from a smooth gradient in the mesiotemporal lobe to higher waviness in the prefrontal cortex (**Figure 3B**). Indeed, subregion smoothness (r_avg_=0.51, p_avg_=0.09) and waviness (r_avg_=0.74, p_avg_=0.011) were correlated between the datasets. Furthermore, in line with our primary analyses, communication efficiency between DMN subregions and the rest of the cortex was higher towards one end of the microstructural axis (r_avg_=-0.38, p_avg-spin_=0.015). This effect was especially pronounced with regards to communication to perceptually-coupled cortical types (koniocortical/eulaminate-III: r_avg_=-0.40/0.37, p_avg-spin_=0.044/0.089). Finally, functional input also tended to decrease along the microstructural axes (r_avg_=-0.26, p_avg-spin_=0.101). Together, these individual-level analyses indicate that the microarchitectural axis of the DMN discriminates a zone of multi-modal convergence from a core that is relatively insulated from external input (**Figure 3C**).

## Discussion

Historically, anatomical details of brain systems have helped to constrain accounts of their function^48,85^. Our study extended this perspective to the default mode network (DMN), one of the most extensively studied yet least well understood systems in the human brain. Leveraging *post-mortem* histology and *in vivo* MRI, we observed pronounced cytoarchitectural heterogeneity within the DMN, showing that the network encompasses types of microarchitecture variably specialised for modality-specific, heteromodal, and self-generated processing^31,43,86^. By combining cytoarchitectural information with structural and functional connectivity, we found that the DMN contains convergence zones that receive input from other cortical regions, as well as a core that is relatively insulated from input. Finally, we found that, unlike other functional networks, outgoing signals of the DMN are of similar strength to different cortical types, meaning the network may be uniquely capable of influencing function across all levels of sensory hierarchies in a relatively coherent manner.

### The DMN harbours a complex landscape of cytoarchitecture and connectivity

Complementary theory- and data-driven analyses revealed the heterogeneous cytoarchitecture of the DMN. On the one hand, comparison of functional and cytoarchitectural atlases showed that the DMN contains a wide range of cortical types, from eulaminate-III to agranular. This type-based analysis demonstrates the extent of cytoarchitectural variation within of the DMN and that it spans multiple steps of laminar elaboration^31,54,87^. On the other hand, applying non-linear dimensionality reduction techniques to an ultrahigh resolution histological reconstruction of a human brain highlighted an axis of cytoarchitectural differentiation, E1, within the DMN that is distinct to the gradient of laminar elaboration. Both the type-based and data-driven axes stretch between the primary sensory areas and the allocortex, but they capture different aspects of cytoarchitectural similarity in eulaminate-II-I and dysgranular cortex. For instance, while cortical types are related combination of qualitative and quantitative measures across cortical layers, the most prominent differences pertain to neuronal density in layers II/III^88^. In contrast, the first data-driven axis is primarily related to cytoarchitectural markers in the mid-to-deep cortical layers. Higher order components, such as E4 and E5, may rather better reflect the cytoarchitectural features captured by cortical types. In addition, cortical types are defined by topology, that is their spatial relations, whereas the data-driven axis is derived in a manner that is agnostic to spatial constraints. The latter approach revealed pronounced cytoarchitectural variation within the DMN that is not as constrained by cortex wide gradients, but rather involves a complex pattern of subregion-specific cytoarchitectural topographies, including both local gradients and interdigitation.

A core principle of neuroanatomy holds that topographies of cortical microstructure, connectivity, and function are intrinsically related^44,61,74,75^. We found a clear example of this relationship in the DMN, whereby the principal cytoarchitectural axis captures differences in structural and functional connectivity to other cortical territories. By combining diffusion-based tractography with physical distance measurements into a model of navigation efficiency^51,76^, we found that the strength of communication between the DMN and other cortical areas was related to the cytoarchitecture of each endpoint. Specifically, regions of the DMN low on E1 exhibited preferentially higher navigation efficiency to granular cortical types. Tract-tracing studies in macaques focusing on circumscribed regions of the DMN, such as the precuneus/posterior cingulate, have shown similar patterns of differential connectivity to primary sensory areas^89,90^. The influence of E1, rather than cortical types, in our analyses, suggests that unique principles of cortical organisation may apply specifically to inter-network connectivity of the DMN.

Repeating the analysis with a regression dynamic causal model (rDCM) of whole-brain effective connectivity^52^, we observed decreasing afferent connectivity along the principal cytoarchitectural axis E1. Areas of the DMN with high afferent connectivity, such as the precuneus and inferior parietal lobe, likely have more supragranular neurons than areas with low afferent connectivity, such as the anterior cingulate and superior frontal gyrus^91,92^. It is possible, therefore, that regions that act as receivers within the DMN may be especially important in feedforward processing^93,94^. This pattern suggests that preferential navigation efficiency from certain subunits of the DMN to more granular types may relate to the speed or directedness of communication, especially given more granular areas exhibit faster intrinsic timescales^95–98^ and sensory areas require high-fidelity information^43^. In contrast, parcels of the DMN with flatter profiles (*i.e.,* higher E1) are more insulated from primary sensory areas [also evident in ^25^] and receive less input from non-DMN cortex. This suggests that the characterisation of the DMN as distant from input^16^ is especially true for those insulated subsections of the DMN (*e.g.,* the anterior cingulate). The degree of insulation may be concordant with suppression during externally-oriented tasks, which is also regionally variable within the DMN^99^. In line with our results, subunits of the DMN high on E1, such as the medial prefrontal cortex, are suppressed for longer than those lower on E1, such as the temporoparietal junction. Taken together the connectivity analyses, therefore, illustrate the complementary functional roles of cytoarchitecturally distinct subunits of the DMN, from receivers on one-side of the cytoarchitectural axis to insulated subunits on the other side.

### Translation from *post-mortem* to *in vivo* research

Our main analyses combined *post-mortem* histology from one individual with in vivo imaging in different populations of healthy individuals. As such, structure-function relationships may be influenced by cross-modal registration as well as inter-individual differences. In this regard, our replication analysis using 7 Tesla MRI shows that fine-grained insights into microarchitecture, connectivity, and function persist at an individual-level and are observable *in vivo.* In other words, they can be seen using microstructural and functional data in a single subject and not just based on population level imaging data or singular *post mortem* resources. We also observed differences between the histological and MRI axes and these may be related to multiple factors including modality (cyto- vs myeloarchitecture), tissue type (*post-mortem* vs *in vivo*) or inter-individual variation. Further work with multiple modalities acquired in a single brain (e.g. MRI and histology or cyto- and myelo-staining) is necessary to determine the source of these differences. Importantly, extending these methods to *in vivo* imaging opens unprecedented possibilities to formally test anatomically grounded hypotheses of the DMN’s role in cognition and behaviour. For example, the present multi-modal model of the DMN, could be combined with psychometric data and experience sampling^17,100,101^ to test how changes in the DMN impact cognitive performance, thought processes and action. Such modelling is a critical next step in evaluating the causal role of the DMN in the brain, as well as the source of its co-fluctuations (for example, by studying the role of neuromodulatory systems^102^).

### The DMN and Cortical Hierarchies

Our investigation of DMN microarchitecture can also help to discern the network’s relationship to cortical hierarchies. Established by foundational research in non-human animals and increasingly confirmed in the human brain, hierarchies are a recurring motif in cortical organisation^39,46^. In general, hierarchical architectures are related to inter-regional variations in temporal dynamics^95,98^ and neural representations. Hierarchies in sensory cortex are well-documented^48^, in part because their properties can be confirmed directly through the stimulation of sensory systems. Hierarchies in association networks, on the other hand, are more challenging to determine^49^, in part due to difficulties in determining a ground truth for their ‘bottom’ and ‘top’. In lieu of such functional evidence, our microarchitectural findings are important because they show the DMN entails two properties of hierarchies, (i) connectivity organisable by distinct levels and (ii) the existence of an apex that is relatively insulated from external input. Unlike sensory hierarchies, however, which increasingly intersect at upper levels, the internal organisation of the DMN is less constrained by spatial gradients and exhibits more balanced interfacing with multiple levels of sensory systems as well as the limbic system. By expanding the conceptualisation of hierarchies beyond sensory systems, our study helps illuminate the diverse nature of information processing in the brain, which is likely to be important in understanding the mechanisms that underpin the role of the DMN in human cognition and action.

Our conceptualisation of the DMN as an association hierarchy expands upon previous ideas, such as the DMN as the apex of^25^ or as a parallel network to^49^ the sensory-fugal hierarchy. Certain features of these theories are concordant with our results, such as (parts of) the DMN being insulated from input and the distinctiveness of information processing in the DMN. However, our analyses demonstrate that connectivity is organised along the most prominent cytoarchitectural axis of the DMN, which is neither nested within or parallel to the sensory-fugal hierarchy. Instead, the DMN seems to protrude from the sensory-fugal hierarchy, with strong afferent connectivity on one end and insulation on the other. The areas with convergent afferents, as well as connections within the DMN, may enable the recombination of neural processes that would not be possible within sensory-fugal processing streams^48^. Such topological complexity is thought to be an important trade-off in development and evolution of biological neural networks^103^ and illustrates how the DMN can play a distinctive role in information integration as an association hierarchy.

### Understanding the role of the DMN in cognition and action

We close by speculating on how our analysis can constrain accounts of the contribution that the DMN makes to human cognition and action. Our study suggests several anatomically grounded hypotheses on how the DMN contributes to a broad range of cognitive states^7,20,58,58^. For instance, the topography of cytoarchitecture can shed light on the different forms of information integration, because more than 90% of cortico-cortical connections are between neighbouring microcircuits^67^. We observed microarchitectural gradients in the mesiotemporal subregion, a pattern previously linked to sequential transformation of signals from low-to higher-order representations^25,104^ and a gradual shift in functional connectivity from the “multiple demand” network to fronto-temporal pole areas^35,105^. In contrast, the interwoven layout of different types of microarchitecture within prefrontal subregions, perhaps related to interdigitation of connections^36,71^, may provide a structural substrate to support domain specialisation^37,38,106^ and cross-domain integration^36^. Understanding the complex cytoarchitectural topography of the prefrontal cortex may also help to understand the region’s functional diversity, which involves both subregional specialisation and functions that are “greater than the sum of its parts”^107^. The presence of both graded and interdigitated motifs within the DMN suggests that when these regions function as a collective, they could contribute to whole brain function in a manner that combines two different types of integration. Furthermore, associations between external and internal modes of cognition and the DMN may be explained by shifting the functional balance from input-oriented to more insulated regions. Such a mechanism would also align with functional imaging studies showing regional differentiation within the DMN for different tasks^58,108^, such as reading *vs.* mind-wandering^109^, which in turn could be linked to how different regions of the DMN participate in or cross-talk with other functional networks^14,110^. Furthermore, in light of the dynamic reconfiguration of functional networks across cognitive states^111^, it will be important to extend the present analysis approach to study the structural properties of the DMN across multiple functional contexts. Additionally, the unique balance that the DMN strikes in terms of its functional output across cortical types may help to unify neural activity across brain systems or verify predictions of the world against memory in real time^20,112^.

Taken together, our study offers a set of anatomical hypotheses on how the human brain may enable the formation of abstract representations and uses these to inform cognition across a range of domains. Specifically, the functional multiplicity of the DMN is pillared upon its internal heterogeneity, possession of receivers and more insulated subunits as well as its balanced communication with all levels of sensory hierarchies. This set of unique features outlines an anatomical landscape within the DMN that may explain why the DMN is involved in states that cross traditional psychological categories and that can have opposing features.

## Concluding Remarks

Since its conceptualisation, the DMN has been marked by controversy. Various approaches produce the DMN, which has led to a certain ontological capaciousness, that is, there is a degree of blurriness about what the DMN is and how to define it^113^. Our study suggests that blurriness of the DMN in both spatial and conceptual terms may be explained by variation in microstructure within sub-regions and their unique connectivity to other regions of cortex. Specifically, the DMN may take on different forms of cognition by recruiting different parts of each subregion, while the broader system maintains the ability to broadcast coherent signals to the rest of the brain. Thus, while the distributed pattern of activity that we’ve come to know as the DMN has been historically enigmatic, our neuroanatomical investigation suggests that this may be feature rather than a bug: It is possible that the capacity for a set of distributed functionally diverse brain regions to operate in a coherent manner may be a core feature of how brain function supports the range of different behaviours that we as a species are capable of engaging in.

## Methods

### Histological data

An ultra-high resolution 3D reconstruction of a sliced and cell-body-stained *post-mortem* human brain from a 65-year-old male was obtained from the open-access BigBrain repository on September 1, 2020 [https://bigbrain.loris.ca/main.php;^55^]. The *post-mortem* brain was paraffin-embedded, coronally sliced into 7,400 20μm sections, silver-stained for cell bodies^114^ and digitised. Manual inspection for artefacts (*i.e.,* rips, tears, shears, and stain crystallisation) was followed by automatic repair procedures, involving non-linear alignment to a *post-mortem* MRI of the same individual acquired prior to sectioning, together with intensity normalisation and block averaging^115^. The 3D reconstruction was implemented with a successive coarse-to-fine hierarchical procedure^116^. We downloaded the 3D volume at 100μm resolution, which was the highest resolution available for the whole brain. Computations were performed on inverted images, where intensity reflects greater cellular density and soma size. Geometric meshes approximating the outer and inner cortical interface (*i.e.,* the GM/CSF boundary and the GM/WM boundary) with 163,842 matched vertices per hemisphere were also obtained^117^.

We constructed 50 equivolumetric surfaces between the outer and inner cortical surfaces . The equivolumetric model compensates for cortical folding by varying the Euclidean distance, ρ, between pairs of intracortical surfaces throughout the cortex to preserve the fractional volume between surfaces^119^. ρ was calculated as follows for each surface

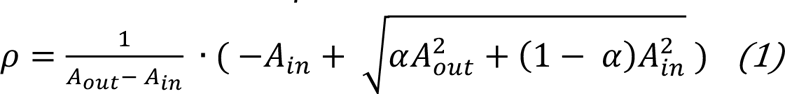

where α represents fraction of the total volume of the segment accounted for by the surface, while A_out_ and A_in_ represent the surface area of the outer and inner cortical surfaces, respectively. Vertex-wise staining intensity profiles were generated by sampling cell-staining intensities along linked vertices from the outer to the inner surface. Smoothing was employed in tangential and axial directions to ameliorate the effects of artefacts, blood vessels, and individual neuronal arrangement. The tangential smoothing across depths was enacted for each staining profile independently, using an iterative piece-wise linear procedure that minimises shrinkage [3 iterations^120^]. Axial surface-wise smoothing was performed at each depth independently and involved moving a 2-vertex FWHM Gaussian kernel across the surface mesh using SurfStat^121^. The staining intensity profiles are made available in the BigBrainWarp toolbox^56^.

### Comparison of cortical atlases

Functional networks were defined using a widely used atlas^2^. The atlas reflects clustering of cortical vertices according to similarity in resting state functional connectivity profiles, acquired in 1000 healthy young adults. Cortical types were assigned to Von Economo areas^54,122^, based on a recent re-analysis of Von Economo micrographs^31^. This classification scheme was used because its criteria are (i) clearly defined, (ii) applied consistently across the entire cortex, (iii) align with Von Economo’s original descriptions and (iv) are supported by multiple histological samples.

Criteria included “development of layer IV, prominence (denser cellularity and larger neurons) of deep (V–VI) or superficial (II–III) layers, definition of sublayers (*e.g.,* IIIa and IIIb), sharpness of boundaries between layers, and presence of large pyramids in superficial layers” ^31^. Thereby, cortical types synopsise degree of granularity, from high laminar elaboration in koniocortical areas, six identifiable layers in Eu-III-I, poorly differentiated layers in dysgranular and absent layers in agranular.

The proportion of DMN vertices assigned to each cortical type was calculated on a common surface template, fsaverage5^123^. The equivalence of cortical type proportions in the DMN and each other functional network was evaluated via pair-wise Kolgomorov-Smirnoff tests. Significant over- or under-representation of each cortical type within the DMN was evaluated with spin permutation testing^124^. Spin permutation testing, used throughout following statistical analyses, involves generating a null distribution by rotating one brain map 10,000 times and recomputing the outcome of interest. Then, we calculate 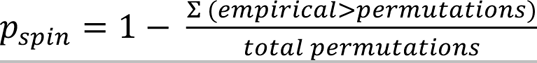 and/or 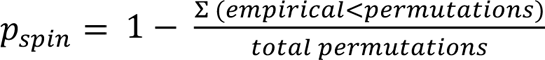. The null distribution preserves the spatial structure of both brain maps, which establishes the plausibility of a random alignment of the maps explaining their statistical correspondence. Generally, we deemed significance p<0.05 for one-tailed tests and p<0.025 for two-tailed tests. Additionally, we used Bonferroni correction when multiple univariate comparisons were made using the same response variable. In the case of the over- or under-representation of specific cortical types within the DMN, we randomly rotated the cortical type atlas, then generated null distributions, representing the number of vertices within the DMN assigned to each type.

The robustness of cytoarchitectural heterogeneity to the DMN definition was assessed with three alternative atlases. Given the origins of the DMN as a “task-negative” set of regions^4,5^, the first alternative atlas involved identifying regions that are consistently deactivated during externally-oriented tasks. In line with a recent review^20^, we used pre-defined contrast maps from 787 healthy young adults of the Human Connectome Project (“HCP_S900_GroupAvg_v1 Dataset”). Each map represents the contrast between BOLD response during a task and at baseline. Fifteen tasks were selected to correspond to early studies of the DMN^5^ [Working Memory (WM)–2 Back, WM-0 Back, WM-Body, WM-Face, WM-Place, WM-Tool, Gambling-Punish, Gambling-Reward, Motor-Average, Social-Random, Social-Theory of Mind, Relational-Match, Relational-Relation, Emotion-Faces, Emotion-Shapes]. For each contrast, task-related deactivation was classed as z-score≤-5, which is consistent with contemporary statistical thresholds used in neuroimaging to reduce false positives^126^. The second alternative atlas represented an independent component analysis of 7,342 task fMRI contrasts. The DMN was specified as the fourth component. The volumetric z-statistic map for that component was projected to the cortical surface for analysis. Thirdly, A probabilistic atlas of the DMN was calculated as the percentage of contrasts with task-related deactivation. The second alternative atlas represented the probability of the DMN at each vertex, calculated across 1029 individual-specific functional network delineations^60^. For each alternative atlas, we calculated the proportions of cortical types across a range of probabilistic thresholds (5-95%, at 5% increments) to determine whether the discovered cytoarchitectural heterogeneity of the DMN was robust to atlas definition.

### Data-driven cytoarchitectural axis within the DMN

The functional network atlas was transformed to the BigBrain surface using a specially optimised multimodal surface matching algorithm^56,64^. The pattern of cytoarchitectural heterogeneity in the DMN was revealed using non-linear manifold learning. The approach involved calculating pair-wise product-moment correlations of BigBrain staining intensity profiles, controlling for the average staining intensity profile within the DMN. Negative values were zeroed to emphasise the non-shared similarities. Diffusion map embedding of the correlation matrix was employed to gain a low dimensional representation of cytoarchitectural patterns^65,124^. Diffusion map embedding belongs to the family of graph Laplacians, which involve constructing a reversible Markov chain on an affinity matrix. Compared to other nonlinear manifold learning techniques, the algorithm is relatively robust to noise and computationally inexpensive^127,128^. A single parameter α controls the influence of the sampling density on the manifold (α = 0, maximal influence; α = 1, no influence). As in previous studies^25,33,124^, we set α = 0.5, a choice retaining the global relations between data points in the embedded space. Notably, different alpha parameters had little to no impact on the first eigenvector (spatial correlation of eigenvectors, r>0.99).

The DMN comprised 71,576 vertices on the BigBrain surface, each associated with approximately 1mm^2^ of surface area. Pair-wise correlation and manifold learning on 71,576 data points was computationally infeasible, however. Thus, we performed a 6-fold mesh decimation on the BigBrain surface to select a subset of vertices that preserve the overall shape of the mesh. Then, we assigned each non-selected vertex to the nearest maintained vertex, determined by shortest path on the mesh (ties were solved by shortest Euclidean distance). Staining intensity profiles were averaged within each surface patch of the DMN, then the dimensionality reduction procedure was employed. Subsequent analyses focused on the first eigenvector (E1), which explained the most variance in the affinity matrix (approximately 28% of variance). Additionally, we repeated this analysis with a highly conservative delineation of the DMN (generated by using the intersection of the three abovementioned alternative atlases), thereby demonstrating that slight variations in atlas definition do not impact the organisation of cytoarchitecture that we discovered in the network. To ensure the spatial pattern depicted by first eigenvector was not purely a product of the selected dimensionality reduction method, we also repeated the procedure using principal component analysis and Laplacian eigenmaps. The first components were near-identical across all approaches (r>0.99).

Local variations in E1 were examined within spatially contiguous subregions of the DMN. Subregions were defined programmatically on the cortical mesh, named according to the gyri they primarily occupy and compared to the Von Economo parcellation (Von Economo areas occupying >10% of the subregion are listed in ascending order in the following brackets); Superior frontal and ACC (FCBm, FB, FA, FDT), middle temporal (TD, PH), inferior parietal (PF, PD, TD), precuneus (PD, LA2, LC1), inferior frontal (FE FDdelta), parahippocampal (HB). Quantitative description of E1 topography within each subregion was achieved with two complementary approaches. First, to characterise the smoothness and complexity of the landscape, we fit polynomial models between E1 and two spatial axes^129^. The spatial axes were derived from an Isomax flattening of each subregion, resulting in a 2D description of each subregion. We compared adjusted R^2^ between subregions within each polynomial order (quadratic, cubic and quartic) using a one-way ANOVA, whereby each subregion was represented by a left and right hemisphere observation. Second, to characterise the bumpiness of subregion landscapes, we adopted an approach from material engineering for characterising the roughness of a surface^69,130^. Specifically, we calculated a waviness metric which reflects the number of intersections of the zero-plane while accounting for the size of the region. As above, we compared waviness between subregions using a one-way ANOVA. Notably, the sensitivity of each approach to variations in E1 topography was validated in a series of simulations, in which we modulated the flatness and bumpiness of the input landscape (**Supplementary Figure 5**).

### MRI acquisition and processing – Primary analyses

Primary MRI analyses were conducted on 40 healthy adults from the microstructure informed connectomics (MICs) cohort (14 females, mean±SD age=30.4±6.7, 2 left-handed)^131^. Scans were completed at the Brain Imaging Centre of the Montreal Neurological Institute and Hospital on a 3T Siemens Magnetom Prisma-Fit equipped with a 64-channel head coil. Two T1w scans with identical parameters were acquired with a 3D-MPRAGE sequence (0.8mm isotropic voxels, TR=2300ms, TE=3.14ms, TI=900ms, flip angle=9°, iPAT=2, matrix=320×320, 224 sagittal slices, partial Fourier=6/8). T1w scans were visually inspected to ensure minimal head motion before they were submitted to further processing. A spin-echo echo-planar imaging sequence with multi-band acceleration was used to obtain DWI data, consisting of three shells with b-values 300, 700, and 2000s/mm^2^ and 10, 40, and 90 diffusion weighting directions per shell, respectively (1.6mm isotropic voxels, TR=3500ms, TE=64.40ms, flip angle=90°, refocusing flip angle=180°, FOV=224×224 mm^2^, slice thickness=1.6mm, multiband factor=3, echo spacing=0.76ms, number of b0 images=3). One 7 min rs-fMRI scan was acquired using multiband accelerated 2D-BOLD echo-planar imaging (3mm isotropic voxels, TR=600ms, TE=30ms, flip angle=52°, FOV=240×240mm^2^, slice thickness=3mm, multiband factor=6, echo spacing=0.54ms). Participants were instructed to keep their eyes open, look at a fixation cross, and not fall asleep. Two spin-echo images with reverse phase encoding were also acquired for distortion correction of the rs-fMRI scans (phase encoding=AP/PA, 3mm isotropic voxels, FOV=240×240mm^2^, slice thickness=3mm, TR=4029 ms, TE=48ms, flip angle=90°, echo spacing=0.54 ms, bandwidth= 2084 Hz/Px).

An open access tool was used for multimodal data processing^132^. Each T1w scan was deobliqued and reoriented. Both scans were then linearly co-registered and averaged, automatically corrected for intensity nonuniformity^133^, and intensity normalized. Resulting images were skull-stripped, and non-isocortical structures were segmented using FSL FIRST^134^. Different tissue types (cortical and subcortical grey matter, white matter, cerebrospinal fluid) were segmented to perform anatomically constrained tractography^135^. Cortical surface segmentations were generated from native T1w scans using FreeSurfer 6.0^123,136,137^. DWI data were pre-processed using MRtrix^138,139^. DWI data underwent b0 intensity normalization, and were corrected for susceptibility distortion, head motion, and eddy currents. Required anatomical features for tractography processing (*e.g.,* tissue type segmentations, parcellations) were non-linearly co-registered to native DWI space using the deformable SyN approach implemented in Advanced Neuroimaging Tools (ANTs)^140^. Diffusion processing and tractography were performed in native DWI space. We performed anatomically-constrained tractography using tissue types segmented from each participant’s pre-processed T1w images registered to native DWI space^135^. We estimated multi-shell and multi-tissue response functions^141^ and performed constrained spherical-deconvolution and intensity normalisation^142^. We initiated the tractogram with 40 million streamlines (maximum tract length=250; fractional anisotropy cutoff=0.06). We applied spherical deconvolution informed filtering of tractograms (SIFT2) to reconstruct whole brain streamlines weighted by cross-sectional multipliers^143^. The reconstructed cross-section streamlines were averaged within 400 spatially contiguous, functionally defined parcels^78^, also warped to DWI space. The rs-fMRI images were pre-processed using AFNI^144^ and FSL^134^. The first five volumes were discarded to ensure magnetic field saturation. Images were reoriented, motion corrected and distortion corrected. Nuisance variable signal was removed using an ICA-FIX classifier^145^ and by performing spike regression. Native timeseries were mapped to individual surface models using a boundary-based registration^146^ and smoothed using a Gaussian kernel (FWHM=10mm, smoothing performed on native midsurface mesh) using workbench^147^. For isocortical regions, timeseries were sampled on native surfaces and averaged within 400 spatially contiguous, functionally defined parcels^78^. For non-isocortical regions, timeseries were averaged within native parcellations of the nucleus accumbens, amygdala, caudate nucleus, hippocampus, pallidum, putamen, and thalamus^134^.

### MRI acquisition and processing – Secondary analyses

Secondary MRI analyses were conducted in 100 unrelated healthy adults (66 females, mean±SD age=28.8±3.8 years) from the minimally preprocessed S900 release of the Human Connectome Project (HCP) . MRI data were acquired on the HCP’s custom 3T Siemens Skyra equipped with a 32-channel head coil. Two T1w images with identical parameters were acquired using a 3D-MPRAGE sequence (0.7mm isotropic voxels, TE=2.14ms, TI=1000ms, flip angle=8°, iPAT=2, matrix=320×320, 256 sagittal slices; TR=2400ms,). Two T2w images were acquired using a 3D T2-SPACE sequence with identical geometry (TR=3200ms, TE=565ms, variable flip angle, iPAT=2). A spin-echo EPI sequence was used to obtain diffusion weighted images, consisting of three shells with *b*-values 1000, 2000, and 3000s/mm^2^ and up to 90 diffusion weighting directions per shell (TR=5520ms, TE=89.5ms, flip angle=78°, refocusing flip angle=160°, FOV=210×180, matrix=178×144, slice thickness=1.25mm, mb factor=3, echo spacing=0.78ms). Four rs-fMRI scans were acquired using multi-band accelerated 2D-BOLD echo-planar imaging (2mm isotropic voxels, TR=720ms, TE=33ms, flip angle=52°, matrix=104×90, 72 sagittal slices, multiband factor=8, 1200 volumes/scan, 3456 seconds). Only the first session was investigated in the present study. Participants were instructed to keep their eyes open, look at a fixation cross, and not fall asleep. Nevertheless, some subjects were drowsy and may have fallen asleep^149^, and the group-averages investigated in the present study do not address these inter-individual differences.

MRI data underwent HCP’s minimal preprocessing^147^. Cortical surface models were constructed using Freesurfer 5.3-HCP^123,136,137^, with minor modifications to incorporate both T1w and T2w^150^. Diffusion MRI data underwent correction for geometric distortions and head motion^147^. Tractographic analysis was based on MRtrix3^138,139^. Response functions for each tissue type were estimated using the dhollander algorithm^151^. Fibre orientation distributions (*i.e.*, the apparent density of fibres as a function of orientation) were modelled from the diffusion-weighted MRI with multi-shell multi-tissue spherical deconvolution^142^, then values were normalised in the log domain to optimise the sum of all tissue compartments towards 1, under constraints of spatial smoothness. Anatomically constrained tractography was performed systematically by generating streamlines using second order integration over fibre orientation distributions with dynamic seeding^143,152^. Streamline generation was aborted when 40 million streamlines had been accepted. We applied spherical deconvolution informed filtering of tractograms (SIFT2) to reconstruct whole brain streamlines weighted by cross-sectional multipliers. The reconstructed cross-section streamlines were averaged within 400 spatially contiguous, functionally defined parcels^78^, also warped to DWI space. The rs-fMRI timeseries were corrected for gradient nonlinearity, head motion, bias field and scanner drifts, then structured noise components were removed using ICA-FIX, further reducing the influence of motion, non-neuronal physiology, scanner artefacts and other nuisance sources^145^. The rs-fMRI data were resampled from volume to MSMAll functionally aligned surface space^153,154^ and averaged within 400 spatially contiguous, functionally defined parcels^78^.

### Modelling structural connectivity with navigation efficiency

Connectivity of DMN subunits was mapped using structural connectomes, derived from diffusion-based tractography. Edge weights of the structural connectomes, representing number of streamlines, were remapped using a log-based transformation: [ −log10(W/(max(W) + min(W>0))]. This log-based transformation attenuates extreme weights and ensures the maximum edge weight is mapped to a positive value. Euclidean distances were calculated between the centroid coordinate of each parcel. Communication in the structural connectome was modelled using navigation^76^, also known as greedy routing^155^. Navigation combines the structural connectome with physical distances, providing a routing strategy that recapitulates invasive, tract-tracing measures of communication^51^. In brief, navigation involves step-wise progression from source node to target node, where each step is determined by spatial proximity to target node.

Specifically, the next node in the path is the neighbour of the current node (*i.e.,* sharing a structural connection) that is closest to the final target node. Navigation is the sum distances of the selected path and navigation efficiency (E_nav_) its inverse; providing an intuitive metric of communication efficiency between two regions. Navigation efficiency was calculated within each hemisphere separately, then concatenated for analyses.

By integrating both topological as well as geometric information in the routing strategy, navigation achieves a topological balance between regularity and randomness that is common for “small-world” networks such as the human brain^156^. Thus, the approach addresses distance bias in group-representative structural connectomes^157^. In prior evaluations^51,76^, navigation was found to both promote a resource-efficient distribution of network information traffic and to explain variation in resting-state functional connectivity. Notably, unlike other commonly studied communication strategies in connectomics (e.g., shortest path routing), navigation does not involve global knowledge of network topology during the node-to-node propagation but simply follows a greedy routing strategy that can be implemented locally, supporting its biological plausibility.

### Modelling functional input and output with effective connectivity

The position of the DMN in large-scale cortical dynamics was explored with regression dynamic causal modelling [rDCM;^52^], a scalable generative model of effective connectivity that allows inferences on the directionality of signal flow, openly available as part of the TAPAS software package^158^. The rDCM was implemented using individual rs-fMRI timeseries. Additionally, an extended version of the rDCM was generated with non-isocortical regions, specifically the nucleus accumbens, amygdala, caudate nucleus, hippocampus, pallidum, putamen, and thalamus.

### Influence of cytoarchitecture on connectivity

Each parcel was labelled according to functional network, modal cortical type and, if part of the DMN, average E1 value. Parcel-average E1 values were calculated by transforming the parcellation scheme to the BigBrain surface and averaging within parcel^56,64^. The following analyses were repeated for E_nav_, effective connectivity derived input and effective connectivity derived output.

First, we selected DMN rows and non-DMN columns of the connectivity matrix. Then, we performed product-moment correlations between E1 and average connectivity to assess the association of the cytoarchitectural axis with connectivity. Next, we stratified the non-DMN columns by cortical type, averaged within type and calculated product-moment correlation between type-average connectivity and E1, providing more specific insight into the relation of the cytoarchitectural axis with connectivity of certain cortical types. For each modality, the correlations were compared to 10,000 spin permutations. P-values were Bonferroni corrected for seven comparisons, resulting in significance threshold of p<0.004 (two-sided test with alpha value of 0.05).

Finally, we estimated the imbalance in connectivity to each cortical type by calculating average connectivity to each type, then calculating the Kullback–Leibler (KL) divergence from a null model with equal average connectivity to each type. The imbalance analysis was repeated for each functional network. In each case, only inter-network connections were included in the calculations. For each modality and each network, we tested whether the KL divergence value was lower than 10,000 spin permutations. P-values were Bonferroni corrected for seven comparisons, resulting in significance threshold of p<0.007 (one-sided test with alpha value of 0.05).

### Individual-level replication with high-field MRI

In the replication, we sought to address two key limitations of the primary analyses. First, due to the unique nature of the BigBrain dataset, cytoarchitectural mapping was based on a single individual, limiting our knowledge of the generalisability of the discovered patterns. Secondly, structural and functional connectivity measurements represented population-averages, thus we were not able to conclude whether the discovered correspondences between cytoarchitecture and connectivity are evident within an individual. To overcome these limitations, we sought to replicate key findings at an individual-level using high-resolution, ultrahigh-field MRI.

Individual-level replication analyses were conducted on 8 healthy adults (5 females, mean±SD age=28±6.3, 1 left-handed). Scans were completed at the Brain Imaging Centre of the Montreal Neurological Institute and Hospital on a 7T Siemens Magnetom Terra System equipped with a 32/8 channel receive/transmit head coil. Two qT1 scans were acquired across two scanning sessions with identical 3D-MP2RAGE sequences (0.5mm isotropic voxels, TR=5170ms, TE=2.44ms, T1_1/2_=1000/3200ms, flip angles=4°, matrix=488×488, slice thickness=0.5mm, partial Fourier=0.75). qT1 maps from the second session were linearly registered to the qT1 maps from the first session, then averaged, to enhanced the signal to noise ratio. A spin-echo echo-planar imaging sequence with multi-band acceleration was used to obtain DWI data, consisting of three shells with b-values 300, 700, and 2000s/mm^2^ and 10, 40, and 90 diffusion weighting directions per shell, respectively (1.1mm isotropic voxels, TR=7383ms, TE=70.6ms, flip angle=90°, matrix=192×192, slice thickness=1.1mm, multiband factor=2, echo spacing=0.26ms, number of b0 images=3, partial Fourier=0.75). One 6 min rs-fMRI scan was acquired using multi-echo, multiband accelerated 2D-BOLD echo-planar imaging (1.9mm isotropic voxels, TR=1690ms, TE_1/2/3_=10.8/27.3/43.8ms, flip angle=67°, matrix=118x118, multiband factor=3, echo spacing=0.54ms, partial Fourier=0.75). Participants were instructed to keep their eyes open, look at a fixation cross, and not fall asleep. Two multiband accelerated spin-echo images with reverse phase encoding were also acquired for distortion correction of the rs-fMRI scans.

The 7T dataset was processed in the same manner as the primary MRI dataset, with two exceptions. qT1 maps were used, rather than T1w images, to construct cortical surfaces, and nuisance variable signal was removed from rs-fMRI using an approach that is specially tailored to multi-echo fMRI (“tedana”)^159^, instead of ICA-FIX, which is optimised for single-echo data. Subsequently, we extracted intracortical profiles from qT1 volumes and determined the principal eigenvector of microstructural differentiation (E1) for each individual using the same procedure as for the histological data. In addition, we used the pre-processed resting state timeseries to produce individual-specific parcellations for each subject, via a pre-trained hierarchical Bayesian model^84^. We subsequently used these parcellations to obtain individual-specific DMNs.

The replication focused on three key results from the primary analysis: (i) DMN subregions differ in terms of the topography of microarchitectural differentiation, which is evident in the roughness of E1. In particular, subregions vary from a gradient in the mesiotemporal lobe to a fluctuating landscape in the prefrontal cortex. (ii) Navigation efficiency decreases along E1, and this effect is especially pronounced for perceptually-coupled cortical types (koniocortical and Eu-III). (iii) Functional input decreases along E1. For each result, we compared statistical outcomes of the primary analysis, derived from BigBrain and population-average connectivity, with individual-level statistical outcomes, derived from the 7T dataset, using product-moment correlations. We report rho and p-values averaged across individuals.

## Data availability

All data that support the findings of this study are openly available. BigBrain is available with LORIS (https://bigbrain.loris.ca/main.php) with preprocessed BigBrain data available in through the BigBrainWarp GitHub repository (https://github.com/caseypaquola/BigBrainWarp). The MICS dataset is available with CONP Portal (https://portal.conp.ca/dataset?id=projects/mica-mics) and the HCP dataset is available with Connectome DB (https://db.humanconnectome.org/).

## Code availability

Custom code for this study, as well as data necessary for reproduction, are openly available on GitHub (https://github.com/caseypaquola/DMN).

## References

1. Power, J. D. et al. Functional network organization of the human brain. Neuron 72, 665–78 (2011).

2. Yeo, B. T. T. et al. The organization of the human cerebral cortex estimated by intrinsic functional connectivity. J. Neurophysiol. 106, 1125–1165 (2011).

3. Greicius, M. D., Krasnow, B., Reiss, A. L. & Menon, V. Functional connectivity in the resting brain: A network analysis of the default mode hypothesis. Proc. Natl. Acad. Sci. 100, 253–258 (2003).

4. Shulman, G. L. et al. Common Blood Flow Changes across Visual Tasks: II. Decreases in Cerebral Cortex. J. Cogn. Neurosci. 9, 648–663 (1997).

5. Raichle, M. E. et al. A default mode of brain function. Proc. Natl. Acad. Sci. U. S. A. 98, 676–682 (2001).

6. Raichle, M. E. The brain’s default mode network. Annu Rev Neurosci 38, 433–447 (2015).

7. Buckner, R. L. & DiNicola, L. M. The brain’s default network: updated anatomy, physiology and evolving insights. Nat. Rev. Neurosci. 20, 593–608 (2019).

8. Simony, E. et al. Dynamic reconfiguration of the default mode network during narrative comprehension. Nat. Commun. 7, 12141–12141 (2016).

9. Yeshurun, Y., Nguyen, M. & Hasson, U. Amplification of local changes along the timescale processing hierarchy. Proc. Natl. Acad. Sci. U. S. A. 114, 9475–9480 (2017).

10. Vatansever, D., Menon, D. K. & Stamatakis, E. A. Default mode contributions to automated information processing. Proc. Natl. Acad. Sci. U. S. A. 114, 12821–12826 (2017).

11. Lanzoni, L. et al. The role of default mode network in semantic cue integration. NeuroImage 219, 117019–117019 (2020).

12. Murphy, C. et al. Modes of operation: A topographic neural gradient supporting stimulus dependent and independent cognition. NeuroImage 186, 487–496 (2019).

13. Braga, R. M., Sharp, D. J., Leeson, C., Wise, R. J. S. & Leech, R. Echoes of the brain within default mode, association, and heteromodal cortices. J. Neurosci. Off. J. Soc. Neurosci. 33, 14031–9 (2013).

14. Spreng, R. N., Stevens, W. D., Chamberlain, J. P., Gilmore, A. W. & Schacter, D. L. Default network activity, coupled with the frontoparietal control network, supports goal-directed cognition. NeuroImage 53, 303–317 (2010).

15. Finc, K. et al. Dynamic reconfiguration of functional brain networks during working memory training. Nat. Commun. 11, 2435 (2020).

16. Murphy, C. et al. Distant from input: Evidence of regions within the default mode network supporting perceptually-decoupled and conceptually-guided cognition. NeuroImage 171, 393–401 (2018).

17. Christoff, K., Gordon, A. M., Smallwood, J., Smith, R. & Schooler, J. W. Experience sampling during fMRI reveals default network and executive system contributions to mind wandering. Proc. Natl. Acad. Sci. U. S. A. 106, 8719–8724 (2009).

18. Karapanagiotidis, T., Bernhardt, B. C., Jefferies, E. & Smallwood, J. Tracking thoughts: Exploring the neural architecture of mental time travel during mind-wandering. NeuroImage 147, 272–281 (2017).

19. Mason, M. F. et al. Wandering minds: The default network and stimulus-independent thought. Science 315, 393–395 (2007).

20. Smallwood, J. et al. The default mode network in cognition: a topographical perspective. Nat. Rev. Neurosci. 2021 228 22, 503–513 (2021).

21. Paquola, C., Amunts, K., Evans, A., Smallwood, J. & Bernhardt, B. Closing the mechanistic gap: the value of microarchitecture in understanding cognitive networks. Trends Cogn. Sci. (2022) doi:10.1016/j.tics.2022.07.001.

22. Alves, P. N. et al. An improved neuroanatomical model of the default-mode network reconciles previous neuroimaging and neuropathological findings. *Commun*. Biol. 2, (2019).

23. Mantini, D. et al. Default Mode of Brain Function in Monkeys. J. Neurosci. 31, 12954–12962 (2011).

24. Buckner, R. L. & Margulies, D. S. Macroscale cortical organization and a default-like apex transmodal network in the marmoset monkey. Nat. Commun. 2019 101 10, 1–12 (2019).

25. Margulies, D. S. et al. Situating the default-mode network along a principal gradient of macroscale cortical organization. Proc. Natl. Acad. Sci. 113, 12574–12579 (2016).

26. Brincat, S. L., Siegel, M., Von Nicolai, C. & Miller, E. K. Gradual progression from sensory to task-related processing in cerebral cortex. (2018) doi:10.1073/pnas.1717075115.

27. Hirabayashi, T. & Miyashita, Y. Computational principles of microcircuits for visual object processing in the macaque temporal cortex. Trends Neurosci. 37, 178–187 (2014).

28. Bechtel, W. & Abrahamsen, A. Explanation: A mechanist alternative. Stud. Hist. Philos. Sci. Part C Stud. Hist. Philos. Biol. Biomed. Sci. 36, 421–441 (2005).

29. Greicius, M. D., Supekar, K., Menon, V. & Dougherty, R. F. Resting-state functional connectivity reflects structural connectivity in the default mode network. Cereb. Cortex 19, 72–78 (2009).

30. Kernbach, J. M. et al. Subspecialization within default mode nodes characterized in 10,000 UK Biobank participants. Proc. Natl. Acad. Sci. U. S. A. 115, 12295–12300 (2018).

31. García-Cabezas, M. Á., Hacker, J. L. & Zikopoulos, B. A Protocol for Cortical Type Analysis of the Human Neocortex Applied on Histological Samples, the Atlas of Von Economo and Koskinas, and Magnetic Resonance Imaging. Front. Neuroanat. 14, 576015 (2020).

32. Bailey, P. & von Bonin, G. *The Isocortex of Man*. (University of Illinois Press, Urbana, 1951).

33. Paquola, C. et al. Microstructural and functional gradients are increasingly dissociated in transmodal cortices. PLoS Biol. 17, e3000284–e3000284 (2019).

34. Royer, J. et al. Myeloarchitecture gradients in the human insula: Histological underpinnings and association to intrinsic functional connectivity. NeuroImage 216, 116859–116859 (2020).

35. Paquola, C. et al. Convergence of cortical types and functional motifs in the human mesiotemporal lobe. eLife 9, e60673 (2020).

36. Goldman-Rakic, P. S. & Schwartz, M. L. Interdigitation of contralateral and ipsilateral columnar projections to frontal association cortex in primates. Science 216, 755–7 (1982).

37. Braga, R. M. & Buckner, R. L. Parallel Interdigitated Distributed Networks within the Individual Estimated by Intrinsic Functional Connectivity. Neuron 95, 457–471.e5 (2017).

38. DiNicola, L. M. & Buckner, R. L. Precision estimates of parallel distributed association networks: evidence for domain specialization and implications for evolution and development. Curr. Opin. Behav. Sci. 40, 120–129 (2021).

39. Markov, N. T. et al. Anatomy of hierarchy: Feedforward and feedback pathways in macaque visual cortex. J. Comp. Neurol. 522, 225–259 (2014).

40. Barbas, H. & Rempel-Clower, N. Cortical structure predicts the pattern of corticocortical connections. Cereb. Cortex 7, 635–646 (1997).

41. Sanides, Friedrich. Die Architektonik des menschlichen Stirnhirns zugleich eine Darstellung der Prinzipien seiner Gestaltung als Spiegel der stammgeschichtlichen Differenzierung der Grosshirnrinde. 201 (Springer, Berlin, 1962).

42. Godlove, D. C., Maier, A., Woodman, G. F. & Schall, J. D. Microcircuitry of Agranular Frontal Cortex: Testing the Generality of the Canonical Cortical Microcircuit. J. Neurosci. 34, 5355–5369 (2014).

43. Mesulam, M.-M. From sensation to cognition. Brain 121, 1013–1052 (1998).

44. Barbas, H. Pattern in the laminar origin of corticocortical connections. J. Comp. Neurol. 252, 415– 422 (1986).

45. Hilgetag, C. C., Medalla, M., Beul, S. F. & Barbas, H. The primate connectome in context: Principles of connections of the cortical visual system. NeuroImage 134, 685–702 (2016).

46. Hilgetag, C. C. & Goulas, A. ‘Hierarchy’ in the organization of brain networks. Philos. Trans. R. Soc. B Biol. Sci. 375, 20190319 (2020).

47. Vezoli, J. et al. Cortical hierarchy, dual counterstream architecture and the importance of top-down generative networks. NeuroImage 225, 117479 (2021).

48. Felleman, D. J. & Van Essen, D. C. Distributed hierarchical processing in the primate cerebral cortex. Cereb. Cortex 1, 1–47 (1991).

49. Buckner, R. L. & Krienen, F. M. The evolution of distributed association networks in the human brain. Trends Cogn. Sci. 17, 648–665 (2013).

50. Goldman-Rakic, P. S. Topography of Cognition: Parallel Distributed Networks in Primate Association Cortex. Annu. Rev. Neurosci. 11, 137–156 (1988).

51. Seguin, C., Razi, A. & Zalesky, A. Inferring neural signalling directionality from undirected structural connectomes. Nat. Commun. 10, 1–13 (2019).

52. Frässle, S. et al. Regression DCM for fMRI. NeuroImage 155, 406–421 (2017).

53. Friston, K. J., Kahan, J., Biswal, B. & Razi, A. A DCM for resting state fMRI. NeuroImage 94, 396–407 (2014).

54. Von Economo, C. & Koskinas, G. *Die Cytoarchitektonik Der Hirnrinde Des Erwachsenen Menschen*. (Springer, Berlin, 1925).

55. Amunts, K. et al. BigBrain: An Ultrahigh-Resolution 3D Human Brain Model. Science 340, 1472– 1475 (2013).

56. Paquola, C. et al. The BigBrainWarp toolbox for integration of BigBrain 3D histology with multimodal neuroimaging. eLife 10, e70119 (2021).

57. Spreng, R. N. & Grady, C. L. Patterns of Brain Activity Supporting Autobiographical Memory, Prospection, and Theory of Mind, and Their Relationship to the Default Mode Network. J. Cogn. Neurosci. 22, 1112–1123 (2010).

58. Andrews-Hanna, J. R., Reidler, J. S., Sepulcre, J., Poulin, R. & Buckner, R. L. Functional-Anatomic Fractionation of the Brain’s Default Network. Neuron 65, 550–562 (2010).

59. Smith, S. M. et al. Correspondence of the brain’s functional architecture during activation and rest. Proc. Natl. Acad. Sci. U. S. A. 106, 13040–5 (2009).

60. Kong, R. et al. Spatial Topography of Individual-Specific Cortical Networks Predicts Human Cognition, Personality, and Emotion. Cereb. Cortex 29, 2533–2551 (2019).

61. Pandya, D., Seltzer, B., Petrides, M. & Cipolloni, P. B. Cerebral Cortex: Architecture, Connections and the Dual Origin Concept. (Oxford University Press, 2015). doi:10.1093/med/9780195385151.001.0001.

62. Bernhardt, B. C. & Singer, T. The neural basis of empathy. Annu. Rev. Neurosci. 35, 1–23 (2012).

63. Hilgetag, C. C., Goulas, A. & Changeux, J.-P. A natural cortical axis connecting the outside and inside of the human brain. Netw. Neurosci. 1–10 (2022) doi:10.1162/netn_a_00256.

64. Lewis, L. B. et al. A multimodal surface matching (MSM) surface registration pipeline to bridge atlases across the MNI and the Freesurfer/Human Connectome Project Worlds. in (Virtual, 2020).

65. Coifman, R. R. & Lafon, S. Diffusion maps. Appl. Comput. Harmon. Anal. 21, 5–30 (2006).

66. Cahalane, D. J., Charvet, C. J. & Finlay, B. L. Systematic, balancing gradients in neuron density and number across the primate isocortex. Front. Neuroanat. 6, 28 (2012).

67. Schüz, A. & Braitenberg, V. The Human Cortical White Matter: Quantitative Aspects of Cortico-Cortical Long-Range Connectivity. in Cortical Areas (CRC Press, 2002).

68. Rosen, B. Q. & Halgren, E. An estimation of the absolute number of axons indicates that human cortical areas are sparsely connected. PLOS Biol. 20, e3001575 (2022).

69. Anon. Surface Texture: Surface Roughness, Waviness and Lay. ANSI Stand *B46* 1 (1978).

70. Braak, H. & Braak, E. On areas of transition between entorhinal allocortex and temporal isocortex in the human brain. Normal morphology and lamina-specific pathology in Alzheimer’s disease. Acta Neuropathol. (Berl.) 68, 325–332 (1985).

71. Selemon, L. D. & Goldman-Rakic, P. S. Common Cortical and Subcortical Targets of the Dorsolateral Prefrontal and Posterior Parietal Cortices in the Rhesus Monkey: Evidence for a Distributed Neural Network Subserving Spatially Guided Behavior. J. Neurosci. 8, 4049–4088 (1988).

72. Jones, E. G. & Powell, T. P. S. An anatomical study of converging sensory pathways within the cerebral cortex of the monkey. Brain 93, 793–820 (1970).

73. Binney, R. J., Parker, G. J. M. & Lambon Ralph, M. A. Convergent connectivity and graded specialization in the rostral human temporal lobe as revealed by diffusion-weighted imaging probabilistic tractography. J. Cogn. Neurosci. 24, 1998–2014 (2012).

74. Hilgetag, C. C. & Grant, S. Cytoarchitectural differences are a key determinant of laminar projection origins in the visual cortex. NeuroImage 51, 1006–1017 (2010).

75. Goulas, A., Margulies, D. S., Bezgin, G. & Hilgetag, C. C. The architecture of mammalian cortical connectomes in light of the theory of the dual origin of the cerebral cortex. Cortex 118, 244–261 (2019).

76. Seguin, C., Van Den Heuvel, M. P. & Zalesky, A. Navigation of brain networks. Proc. Natl. Acad. Sci. U. S. A. 115, 6297–6302 (2018).

77. Barbas, H. General Cortical and Special Prefrontal Connections: Principles from Structure to Function. Annu. Rev. Neurosci. 38, 269–289 (2015).

78. Schaefer, A. et al. Local-Global Parcellation of the Human Cerebral Cortex from Intrinsic Functional Connectivity MRI. Cereb. Cortex N. Y. N 1991 28, 3095–3114 (2018).

79. Frässle, S. et al. Regression dynamic causal modeling for resting-state fMRI. Hum. Brain Mapp. hbm.25357-hbm.25357 (2021) doi:10.1002/hbm.25357.

80. Paquola, C. & Hong, S.-J. The potential of myelin-sensitive imaging: Redefining spatiotemporal patterns of myeloarchitecture. Biol. Psychiatry (2022) doi:10.1016/j.biopsych.2022.08.031.

81. Stüber, C. et al. Myelin and iron concentration in the human brain: A quantitative study of MRI contrast. NeuroImage 93, 95–106 (2014).

82. Hellwig, B. How the myelin picture of the human cerebral cortex can be computed from cytoarchitectural data. A bridge between von Economo and Vogt. J. Hirnforsch. 34, 387–402 (1993).

83. Sanides, F. The Cyto-myeloarchitecture of the Human Frontal Lobe and its Relation to Phylogenetic Differentiation of the Cerebral Cortex. J. Für Hirnforsch. 6, 269–282 (1964).

84. Kong, R. et al. Individual-Specific Areal-Level Parcellations Improve Functional Connectivity Prediction of Behavior. Cereb. Cortex 31, 4477–4500 (2021).

85. Chalfie, M. et al. The neural circuit for touch sensitivity in Caenorhabditis elegans. J. Neurosci. Off. J. Soc. Neurosci. 5, 956–964 (1985).

86. García-Cabezas, M. Á., Zikopoulos, B. & Barbas, H. The Structural Model: a theory linking connections, plasticity, pathology, development and evolution of the cerebral cortex. Brain Struct. Funct. 224, 985–1008 (2019).

87. Von Economo, C. & Triarhou, L. C. Cellular Structure of the Human Cerebral Cortex. Cellular Structure of the Human Cerebral Cortex (S. Karger AG, 2009). doi:10.1093/brain/awp268.

88. Dombrowski, S. M. Quantitative Architecture Distinguishes Prefrontal Cortical Systems in the Rhesus Monkey. Cereb. Cortex 11, 975–988 (2001).

89. Kobayashi, Y. & Amaral, D. G. Macaque monkey retrosplenial cortex: II. Cortical afferents. J. Comp. Neurol. 466, 48–79 (2003).

90. Margulies, D. S. et al. Precuneus shares intrinsic functional architecture in humans and monkeys. Proc. Natl. Acad. Sci. U. S. A. 106, 20069–74 (2009).

91. Goulas, A., Majka, P., Rosa, M. G. P. & Hilgetag, C. C. A blueprint of mammalian cortical connectomes. PLoS Biol. 17, e2005346–e2005346 (2019).

92. Pronold, J. et al. Multi-Scale Spiking Network Model of Human Cerebral Cortex. 2023.03.23.533968 Preprint at 10.1101/2023.03.23.533968 (2023).

93. Rockland, K. S. & Pandya, D. N. Laminar origins and terminations of cortical connections of the occipital lobe in the rhesus monkey. Brain Res. 179, 3–20 (1979).

94. Schmidt, M. et al. A multi-scale layer-resolved spiking network model of resting-state dynamics in macaque visual cortical areas. PLOS Comput. Biol. 14, e1006359 (2018).

95. Chaudhuri, R., Knoblauch, K., Gariel, M.-A., Kennedy, H. & Wang, X.-J. A Large-Scale Circuit Mechanism for Hierarchical Dynamical Processing in the Primate Cortex. Neuron 88, 419–431 (2015).

96. Gao, R., Van den Brink, R. L., Pfeffer, T. & Voytek, B. Neuronal timescales are functionally dynamic and shaped by cortical microarchitecture. eLife 9, 1–44 (2020).

97. Ito, T., Hearne, L. J. & Cole, M. W. A cortical hierarchy of localized and distributed processes revealed via dissociation of task activations, connectivity changes, and intrinsic timescales. NeuroImage 221, 117141–117141 (2020).

98. Murray, J. D. et al. A hierarchy of intrinsic timescales across primate cortex. Nat. Neurosci. 17, 1661–3 (2014).

99. Ossandón, T. et al. Transient suppression of broadband gamma power in the default-mode network is correlated with task complexity and subject performance. J. Neurosci. 31, 14521–14530 (2011).

100. De Marco, M. et al. Cognitive stimulation of the default-mode network modulates functional connectivity in healthy aging. Brain Res. Bull. 121, 26–41 (2016).

101. Sormaz, M. et al. Default mode network can support the level of detail in experience during active task states. Proc. Natl. Acad. Sci. U. S. A. 115, 9318–9323 (2018).

102. van den Brink, R. L., Pfeffer, T. & Donner, T. H. Brainstem Modulation of Large-Scale Intrinsic Cortical Activity Correlations. Front. Hum. Neurosci. 13, (2019).

103. Bassett, D. S. et al. Efficient physical embedding of topologically complex information processing networks in brains and computer circuits. PLoS Comput. Biol. 6, e1000748 (2010).

104. Sepulcre, J., Sabuncu, M. R., Yeo, T. B., Liu, H. & Johnson, K. A. Stepwise connectivity of the modal cortex reveals the multimodal organization of the human brain. J. Neurosci. 32, 10649– 10661 (2012).

105. Duncan, J. The multiple-demand (MD) system of the primate brain: mental programs for intelligent behaviour. Trends Cogn. Sci. 14, 172–179 (2010).

106. Braga, R. M., DiNicola, L. M., Becker, H. C. & Buckner, R. L. Situating the left-lateralized language network in the broader organization of multiple specialized large-scale distributed networks. J. Neurophysiol. 124, 1415–1448 (2020).

107. Wilson, C. R. E., Gaffan, D., Browning, P. G. F. & Baxter, M. G. Functional localization within the prefrontal cortex: missing the forest for the trees? Trends Neurosci. 33, 533–40 (2010).

108. Smith, V., Mitchell, D. J. & Duncan, J. Role of the Default Mode Network in Cognitive Transitions. Cereb. Cortex 28, 3685–3696 (2018).

109. Zhang, M. et al. Perceptual coupling and decoupling of the default mode network during mind-wandering and reading. eLife 11, e74011 (2022).

110. Faskowitz, J., Esfahlani, F. Z., Jo, Y., Sporns, O. & Betzel, R. F. Edge-centric functional network representations of human cerebral cortex reveal overlapping system-level architecture. Nat. Neurosci. 23, 1644–1654 (2020).

111. Salehi, M., Karbasi, A., Barron, D. S., Scheinost, D. & Constable, R. T. Individualized functional networks reconfigure with cognitive state. NeuroImage 206, 116233 (2020).

112. Yeshurun, Y., Nguyen, M. & Hasson, U. The default mode network: where the idiosyncratic self meets the shared social world. Nat. Rev. Neurosci. 22, 181–192 (2021).

113. Callard, F. & Margulies, D. S. What we talk about when we talk about the default mode network. Front. Hum. Neurosci. 8, 619 (2014).

114. Merker, B. Silver staining of cell bodies by means of physical development. J. Neurosci. Methods 9, 235–241 (1983).

115. Lepage, C. Y. et al. Automatic Repair of Acquisition Defects in Reconstruction of Histology Sections of a Human Brain. in Annual Meeting of the Organization for Human Brain Mapping (Barcelona, 2010).

116. Mohlberg, H., Tweddell, B., Lippert, T. & Amunts, K. Workflows for Ultra-High Resolution 3D Models of the Human Brain on Massively Parallel Supercomputers. in 15–27 (Springer, Cham, 2016). doi:10.1007/978-3-319-50862-7_2.

117. Lewis, L. B. et al. BigBrain: Initial Tissue Classification and Surface Extraction. in (Hamburg, 2014).

118 Wagstyl, K., Paquola, C., Bethlehem, R. & Huth, A. kwagstyl/surface_tools: Initial release of equivolumetric surfaces 10.5281/ZENODO.1412054 (2018).

119. Waehnert, M. D. et al. Anatomically motivated modeling of cortical laminae. NeuroImage 93, 210– 220 (2014).

120. Taubin, G. Curve and surface smoothing without shrinkage. in IEEE International Conference on Computer Vision 852–857 (IEEE, 1995). doi:10.1109/iccv.1995.466848.

121. Worsley, K., et al. SurfStat: A Matlab toolbox for the statistical analysis of univariate andmultivariate surface and volumetric data using linear mixed effects modelsand random field theory. in Human Brain Mapping (2009).

122. Scholtens, L. H., de Reus, M. A., de Lange, S. C., Schmidt, R. & van den Heuvel, M. P. An MRI Von Economo-Koskinas atlas. NeuroImage (2016) doi:10.1016/j.neuroimage.2016.12.069.

123. Dale, A. M., Fischl, B. & Sereno, M. I. Cortical surface-based analysis. I. Segmentation and surface reconstruction. Neuroimage 9, 179–194 (1999).

124. Vos de Wael, R., et al. BrainSpace: a toolbox for the analysis of macroscale gradients in neuroimaging and connectomics datasets. *Commun*. Biol. 3, 103 (2020).

125. Alexander-Bloch, A. F. et al. On testing for spatial correspondence between maps of human brain structure and function. NeuroImage 178, 540–551 (2018).

126. Eklund, A., Nichols, T. E. & Knutsson, H. Cluster failure: Why fMRI inferences for spatial extent have inflated false-positive rates. Proc Natl Acad Sci U A 113, 7900–7905 (2016).

127. Tenenbaum, J. B., De Silva, V. & Langford, J. C. A global geometric framework for nonlinear dimensionality reduction. Science 290, 2319–23 (2000).

128. Von Luxburg, U. A tutorial on spectral clustering. Stat. Comput. 17, 395–416 (2007).

129. D’Errico, J. polyfitn. (2023)

130. Gadelmawla, E. S., Koura, M. M., Maksoud, T. M. A., Elewa, I. M. & Soliman, H. H. Roughness parameters. J. Mater. Process. Technol. 123, 133–145 (2002).

131. Royer, J., et al. An Open MRI Dataset for Multiscale Neuroscience. bioRxiv (2021) 10.1101/2021.08.04.454795.

132. Cruces, R. R. et al. Micapipe: A pipeline for multimodal neuroimaging and connectome analysis. NeuroImage 263, 119612 (2022).

133. Tustison, N. J. & Avants, B. B. Explicit B-spline regularization in diffeomorphic image registration. *Front*. Neuroinformatics 7, 39–39 (2013).

134. Jenkinson, M., Beckmann, C. F., Behrens, T. E., Woolrich, M. W. & Smith, S. M. Fsl. Neuroimage 62, 782–790 (2012).

135. Smith, R. E., Tournier, J.-D., Calamante, F. & Connelly, A. Anatomically-constrained tractography: improved diffusion MRI streamlines tractography through effective use of anatomical information. NeuroImage 62, 1924–38 (2012).

136. Fischl, B., Sereno, M. I. & Dale, A. M. Cortical surface-based analysis. II: Inflation, flattening, and a surface-based coordinate system. Neuroimage 9, 195–207 (1999).

137. Fischl, B., Sereno, M. I., Tootell, R. B. & Dale, A. M. High-resolution intersubject averaging and a coordinate system for the cortical surface. Hum. Brain Mapp. 8, 272–284 (1999).

138. Tournier, J.-D., Calamante, F. & Connelly, A. MRtrix: diffusion tractography in crossing fiber regions. Int. J. Imaging Syst. Technol. 22, 53–66 (2012).

139. Tournier, J.-D. et al. MRtrix3: A fast, flexible and open software framework for medical image processing and visualisation. NeuroImage 116137–116137 (2019).

140. Avants, B. B., Epstein, C. L., Grossman, M. & Gee, J. C. Symmetric diffeomorphic image registration with cross-correlation: Evaluating automated labeling of elderly and neurodegenerative brain. Med. Image Anal. 12, 26–41 (2008).

141. Christiaens, D. et al. Global tractography of multi-shell diffusion-weighted imaging data using a multi-tissue model. Neuroimage 123, 89–101 (2015).

142. Jeurissen, B., Tournier, J.-D., Dhollander, T., Connelly, A. & Sijbers, J. Multi-tissue constrained spherical deconvolution for improved analysis of multi-shell diffusion MRI data. NeuroImage 103, 411–426 (2014).

143. Smith, R. E., Tournier, J. D., Calamante, F. & Connelly, A. SIFT2: Enabling dense quantitative assessment of brain white matter connectivity using streamlines tractography. NeuroImage 119, 338–351 (2015).

144. Cox, R. W. AFNI: software for analysis and visualization of functional magnetic resonance neuroimages. Comput. Biomed. Res. 29, 162–173 (1996).

145. Salimi-Khorshidi, G. et al. Automatic denoising of functional MRI data: Combining independent component analysis and hierarchical fusion of classifiers. NeuroImage 90, 449–468 (2014).

146. Greve, D. N. & Fischl, B. Accurate and robust brain image alignment using boundary-based registration. NeuroImage 48, 63–72 (2009).

147. Glasser, M. F. et al. The minimal preprocessing pipelines for the Human Connectome Project. NeuroImage 80, 105–124 (2013).

148. Van Essen, D. C. et al. The WU-Minn Human Connectome Project: An overview. NeuroImage 80, 62–79 (2013).

149. Glasser, M. F. et al. Using temporal ICA to selectively remove global noise while preserving global signal in functional MRI data. NeuroImage 181, 692–717 (2018).

150. Glasser, M. F. & Van Essen, D. C. Mapping human cortical areas in vivo based on myelin content as revealed by T1-and T2-weighted MRI. J. Neurosci. Off. J. Soc. Neurosci. 31, 11597–616 (2011).

151. Dhollander, T., Raffelt, D. & Connelly, A. Unsupervised 3-tissue response function estimation from single-shell or multi-shell diffusion MR data without a co-registered T1 image. in ISMRM Workshop on Breaking the Barriers of Diffusion MRI 5–5 (2016).

152. Tournier, J. D., Calamante, F. & Connelly, A. Robust determination of the fibre orientation distribution in diffusion MRI: Non-negativity constrained super-resolved spherical deconvolution. NeuroImage 35, 1459–1472 (2007).

153. Robinson, E. C. et al. MSM: A new flexible framework for multimodal surface matching. NeuroImage 100, 414–426 (2014).

154. Robinson, E. C. et al. Multimodal surface matching with higher-order smoothness constraints. NeuroImage 167, 453–465 (2018).

155. Muscoloni, A., Thomas, J. M., Ciucci, S., Bianconi, G. & Cannistraci, C. V. Machine learning meets complex networks via coalescent embedding in the hyperbolic space. Nat. Commun. 8, 1–19 (2017).

156. Bassett, D. S. & Bullmore, E. Small-World Brain Networks. The Neuroscientist 12, 512–523 (2006).

157. Betzel, R. F., Griffa, A., Hagmann, P. & Mišić, B. Distance-dependent consensus thresholds for generating group-representative structural brain networks. Netw. Neurosci. 3, 475–496 (2019).

158. Frässle, S. et al. TAPAS: An Open-Source Software Package for Translational Neuromodeling and Computational Psychiatry. Front. Psychiatry 12, (2021).

159. DuPre, E. et al. TE-dependent analysis of multi-echo fMRI with *tedana*. J. Open Source Softw. 6, 3669 (2021).

160. Salo, T. et al. NiMARE: Neuroimaging Meta-Analysis Research Environment. NeuroLibre Reprod. Prepr. Serv. 1, 7 (2022).

161. Yarkoni, T., Poldrack, R. A., Nichols, T. E., Van Essen, D. C. & Wager, T. D. Large-scale automated synthesis of human functional neuroimaging data. Nat. Methods 8, 665–670 (2011).

