## Supplementary for "Architecture of the Human Default Mode Network: cytoarchitecture, wiring and signal flow"

### SUPPLEMENTARY INFORMATION

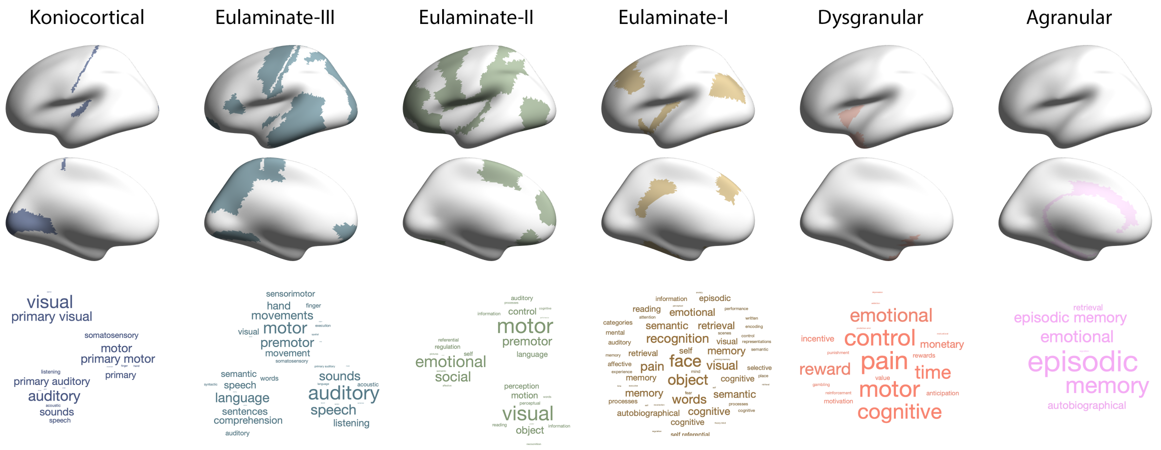


**Supplementary Figure 1:** Meta-analytic functional decoding of the cortical type atlas supports the association, described in literature reviews[43], between the gradient of cortical types and a shift in function from primary sensory to unimodal to heteromodal to memory-related processes. Using meta-analytic maps of thousands of functional MRI[160,161], we extracted terms that were consistently associated with increased activity within the specific cortical type (threshold z-statistic>2). The size of each word reflects the relative strength of its association with the cortical type. Only psychological constructs were retained in the term lists (thus excluding anatomical terms, e.g. "V1", and experiment-related terms, e.g. "healthy controls"). Decoding was performed within spatially contiguous subregions for Kon, Eu-III and Eu-II, because no terms exceeded the threshold when the subregions were combined, due to the distinctive unimodal functions of each subregion.

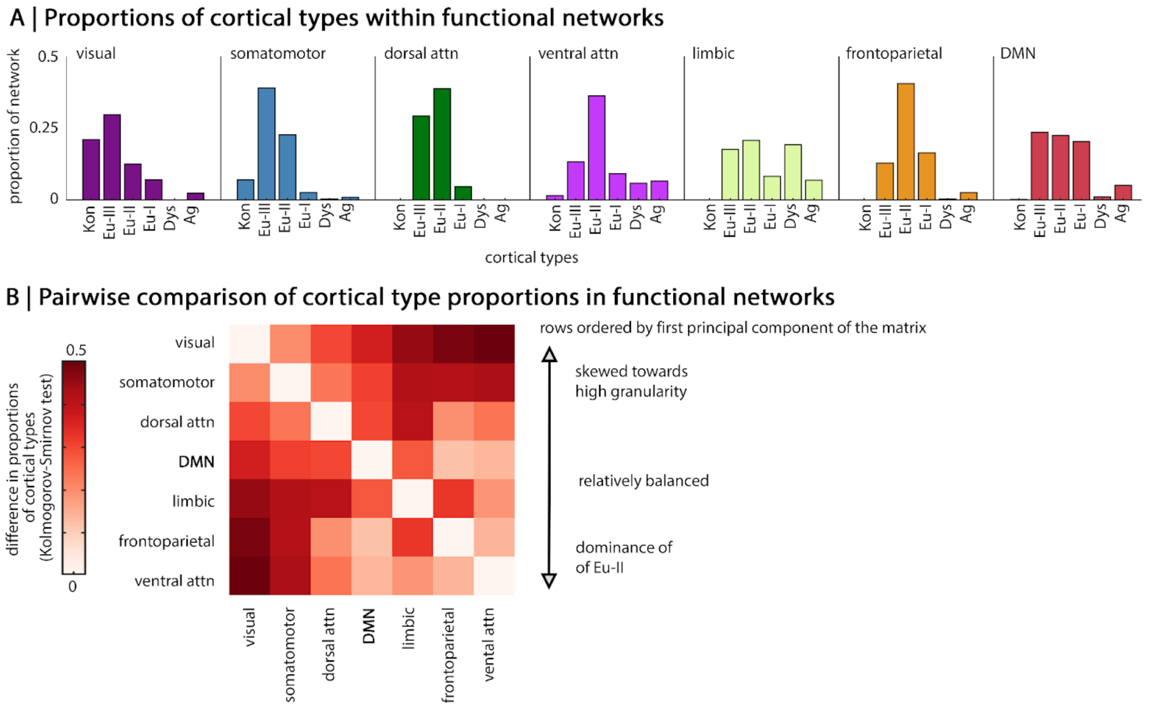


**Supplementary Figure 2: A)** Bar charts illustrate the proportion of cortical types within each functional network **(**for further details, see **Supplementary Table 2. B)** Matrix illustrating the outcome of pair-wise Kolmogorov-Smirnov tests, whereby darker colours reflect greater difference in the cortical type make-up of the functional networks. Rows and columns of the matrix are ordered according to the first principal component, thereby showing that the DMN occupies a middle ground between the functional networks skewed towards high granularity and the functional networks dominated by eulaminate-II.

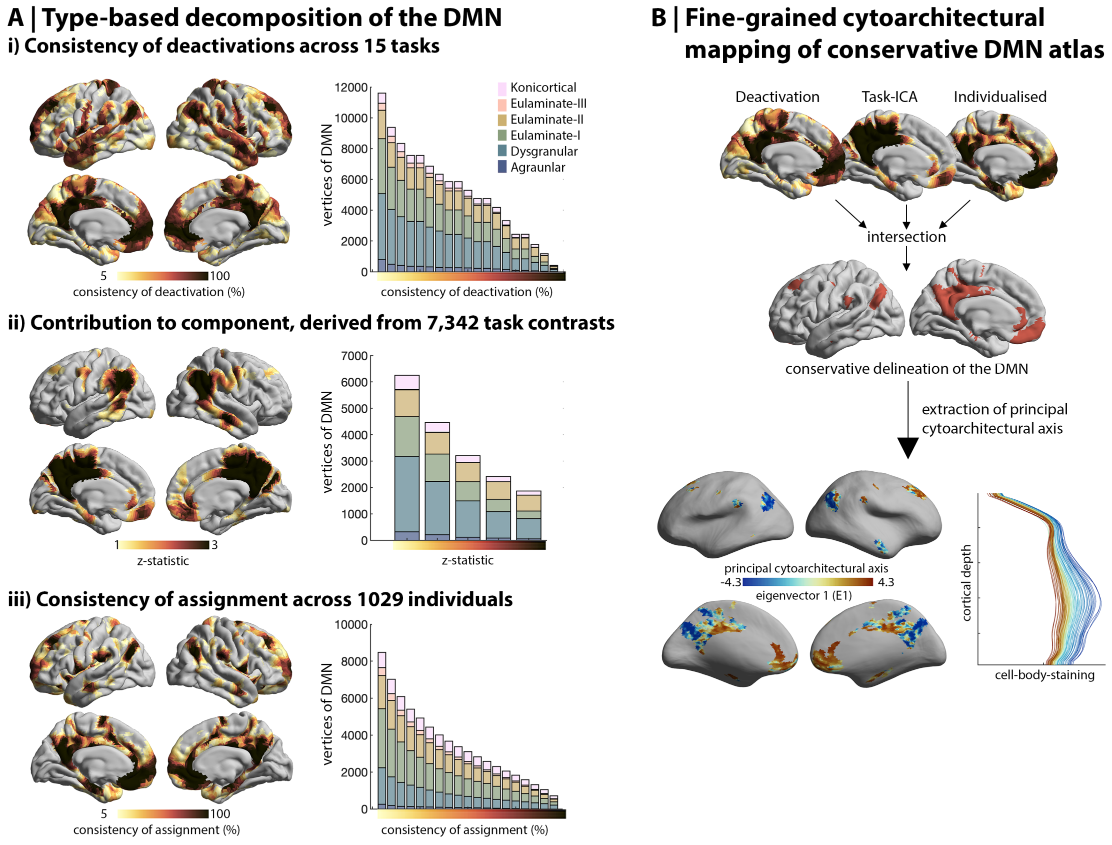


**Supplementary Figure 3: Cytoarchitectural heterogeneity in the DMN replicated with alternative atlases. A)** The diverse cytoarchitectural composition of the DMN was also evident using alternative atlas definitions. Stacked boxplots illustrate the number of vertices assigned to each cortical type within the atlas with increasingly conservative thresholds for inclusion in the DMN represented along the x-axis. **i)** DMN based on consistency of deactivation during perceptually-driven tasks. Vertex-wise change in the BOLD response were calculated across 787 subjects in Human Connectome Project during fifteen perceptually-driven tasks. Surface projections show the consistency of deactivations ($z \leq -5$) across the tasks[20]. **ii)** Association (z-statistic) of each vertex to the DMN derived from an independent component analysis of 7,342 task contrasts[59]. **iii)** Probability of the DMN at each vertex, calculated across 1029 individual-specific functional network delineations[60]. Proportion of the DMN assigned to each cortical type, where the DMN is defined variably based on different consistency thresholds. **B)** Using an intersection of the three approaches in part A, we created a highly conservative delineation of the DMN. Specifically, vertices were included in the conservative atlas if (i) deactivations were observed in more than a quarter of perceptually-driven tasks, (ii) contribution to the task-ICA exceeded a z-statistic of 1 and (iii) assignment to the DMN was observed in more than a quarter of individuals. Subsequently, we replicated the procedure in the primary analysis to extract the principal cytoarchitectural axis. Notably, similar patterns of cytoarchitectural differentiation are evident in this conservative delineation of the DMN. The conservative cytoarchitectural axis also captures a variation from peaked to flat profiles.

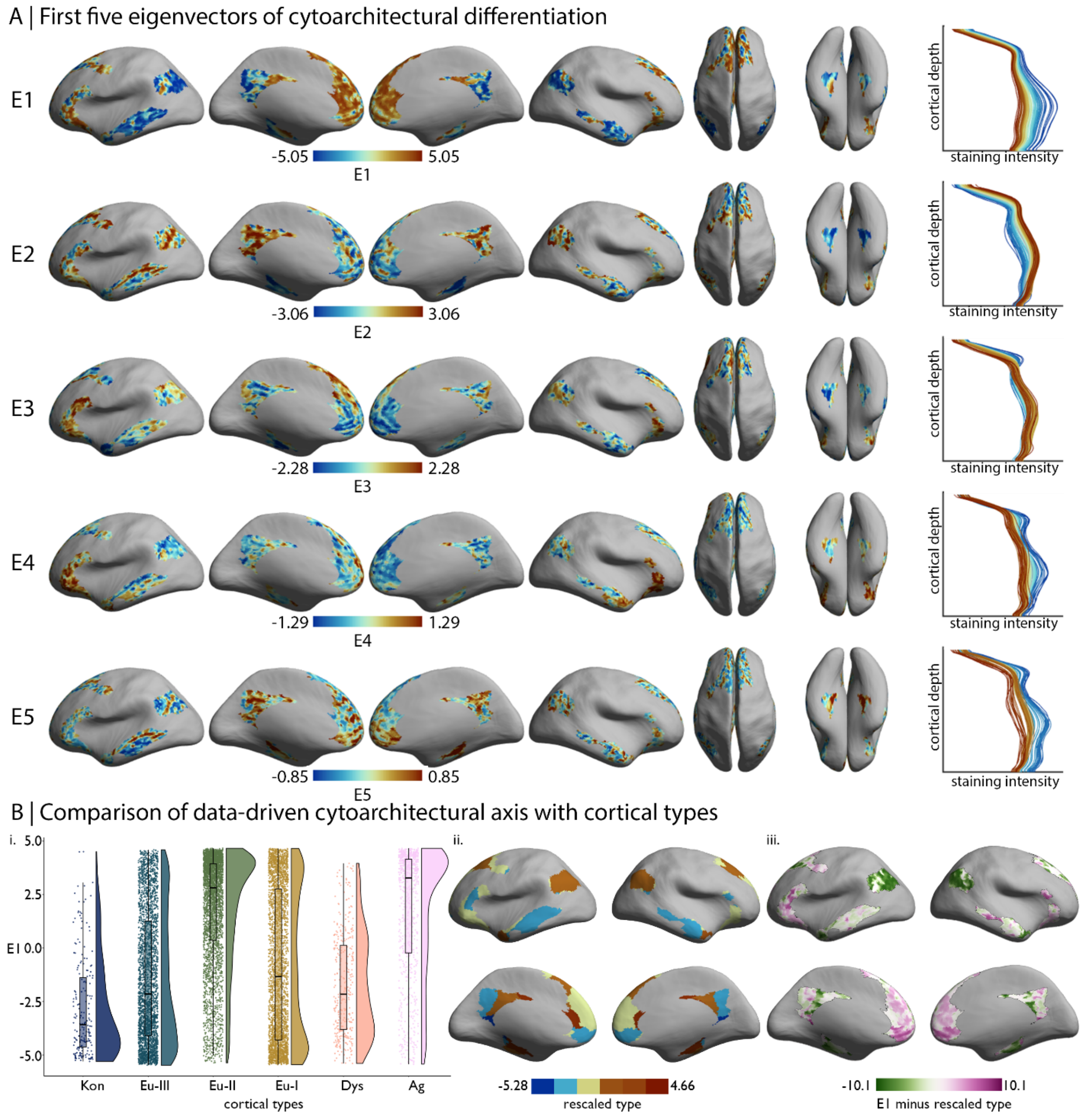


**Supplementary Figure 4: A)** First five eigenvectors projected on the inflated BigBrain surface. For line plots on the right, staining intensity profiles were averaged within 100 bins of the respective eigenvector and coloured by eigenvector position. **B) i.** Raincloud plot shows the distribution of E1 across cortical types. **ii.** Cortical type assignment (1:6) was rescaled to the range of E1 then subtracted from E1, producing a deviation map that highlights where the type-based and data-driven depictions of DMN cytoarchitecture differ. Negative values indicate lower E1 than expected by a linear relationship with cortical type, whereas positive values indicate higher than predicted E1. Thus, the E1 pattern is distinct to the gradient of laminar elaboration that is captured by the cortical types. Both are anchored by koniocortex on one side and agranular cortex on the other, but they differ in the ordering of Eu-and dysgranular areas.

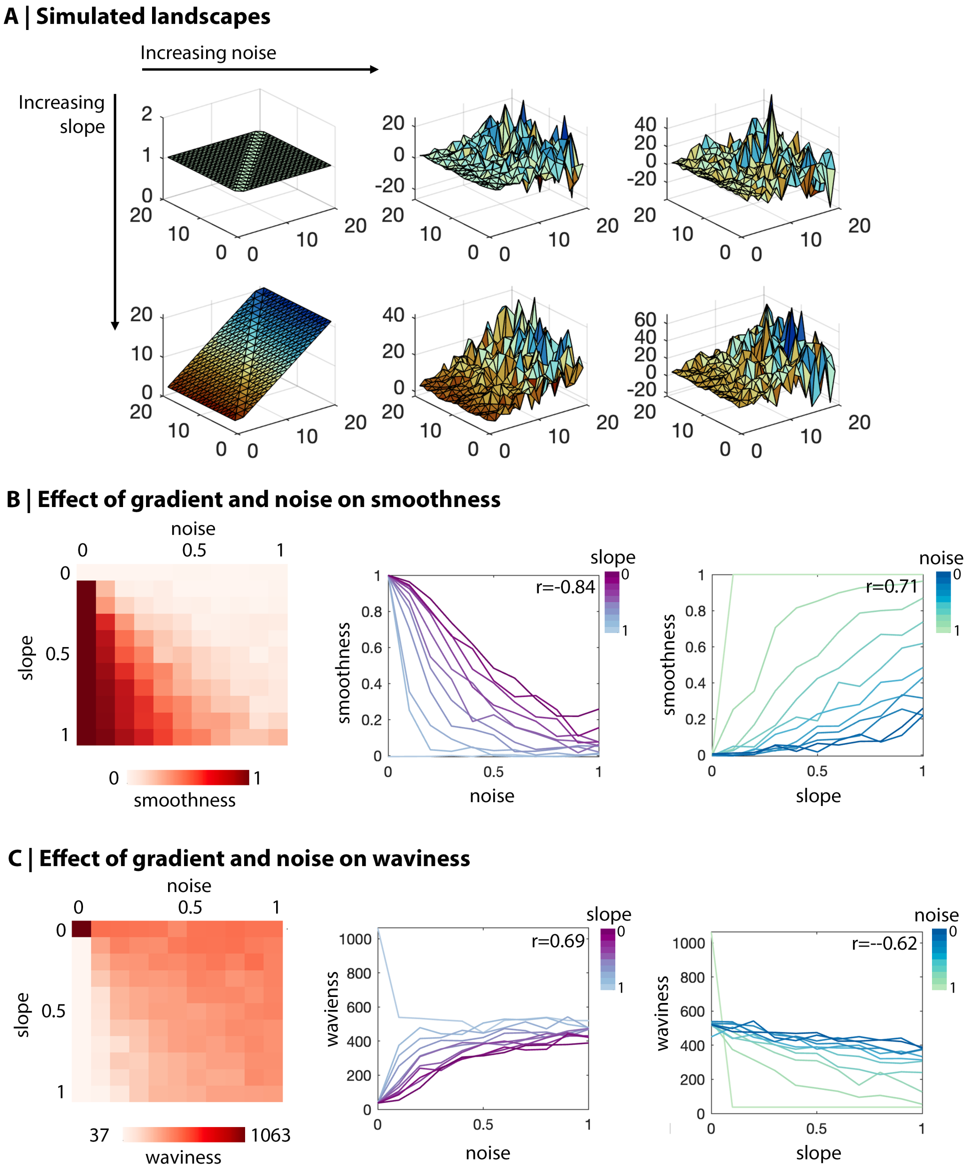


**Supplementary Figure 5:** As expected, smoothness decreases with noise and increases with slope, whereas waviness increases with noise and decreases with slope. **A)** We simulated 121 landscapes with varied slopes and bumpiness (noise). x and y values were identical in all landscapes, while the z-axis – reflecting E1 topography in the main study – was modulated in each simulation. The z-axis value was calculated as "(x * *slope*) + (*rand* * *sigma*)", where *slope* is a value within [0:0.1:1], *rand* is a vector of normally distributed pseudorandom numbers the length of x and *sigma* is the product of x and a value within [0:0.1:1]. **B-C)** ***Left.*** Each square of the matrix represents a simulated landscape, with rows reflecting increasing slope and columns reflecting increasing noise. ***Centre-Right.*** Line plots show the outcome metrics of simulations per row and column, respectively. r-values represent the outcome of partial product-moment correlations (e.g. correlation of smoothness with noise, controlling for slope).

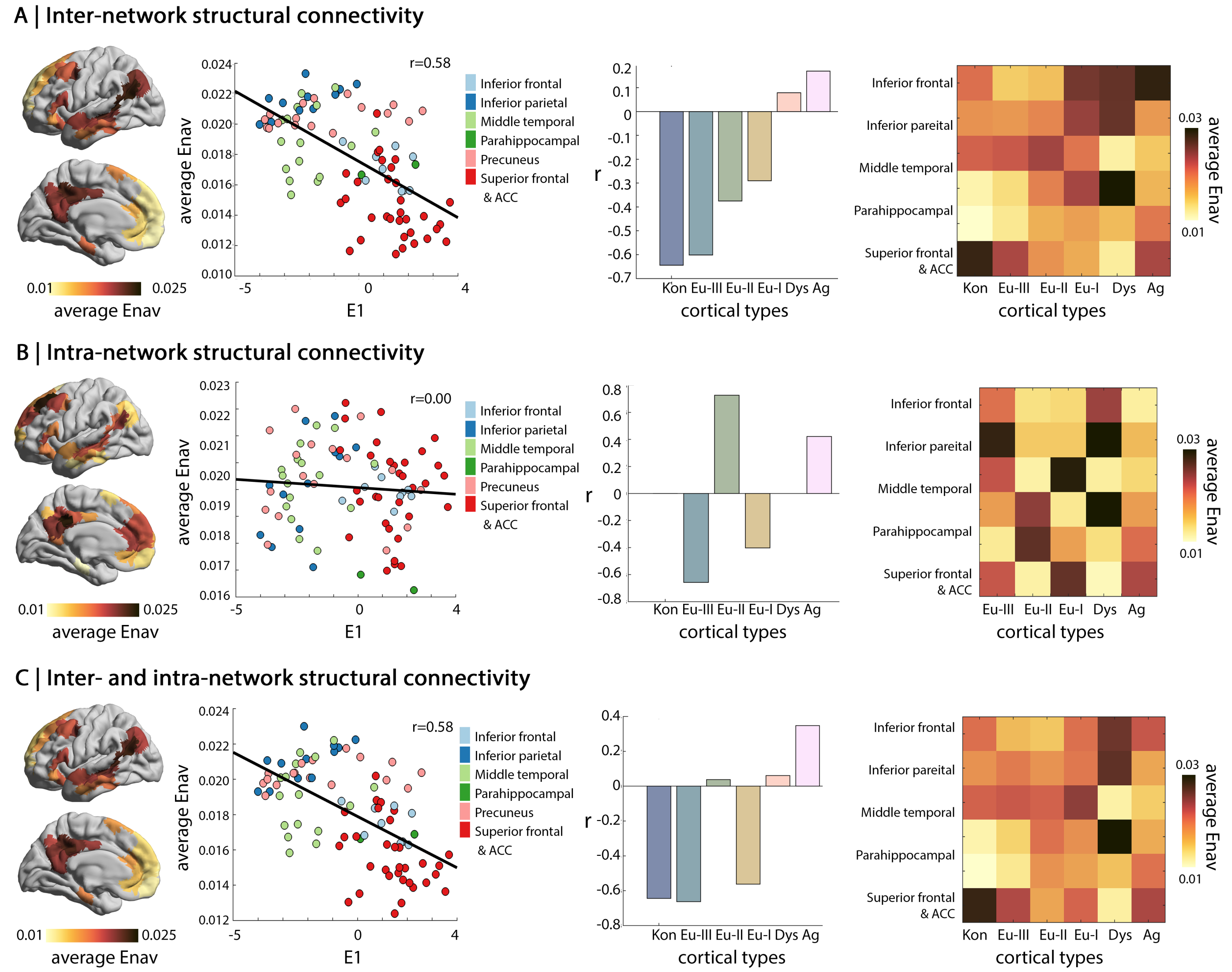


**Supplementary Figure 6:** Variations in navigation efficiency as a function of the cytoarchitectural axis within the DMN, DMN subregion and cortical type. Panel A) involves connections from each node of the DMN with all nodes outside the DMN (as in the primary analysis), Panel B) connections from each node of the DMN to all other nodes of the DMN and Panel C) connections from each node of the DMN to all other nodes. *Far left.* Cortical maps show average navigation efficiency. *Centre left.* Scatterplots show the correlation of the cytoarchitectural axis (E1) with average navigation efficiency, with points coloured by the seed parcel's position within the DMN. *Centre right*. Bar plots show the linear correlation coefficient (r) of E1 with average navigation efficiency to each cortical type. *Far right*. Matrix shows the average navigation efficiency between each subregion of the DMN and each cortical type.

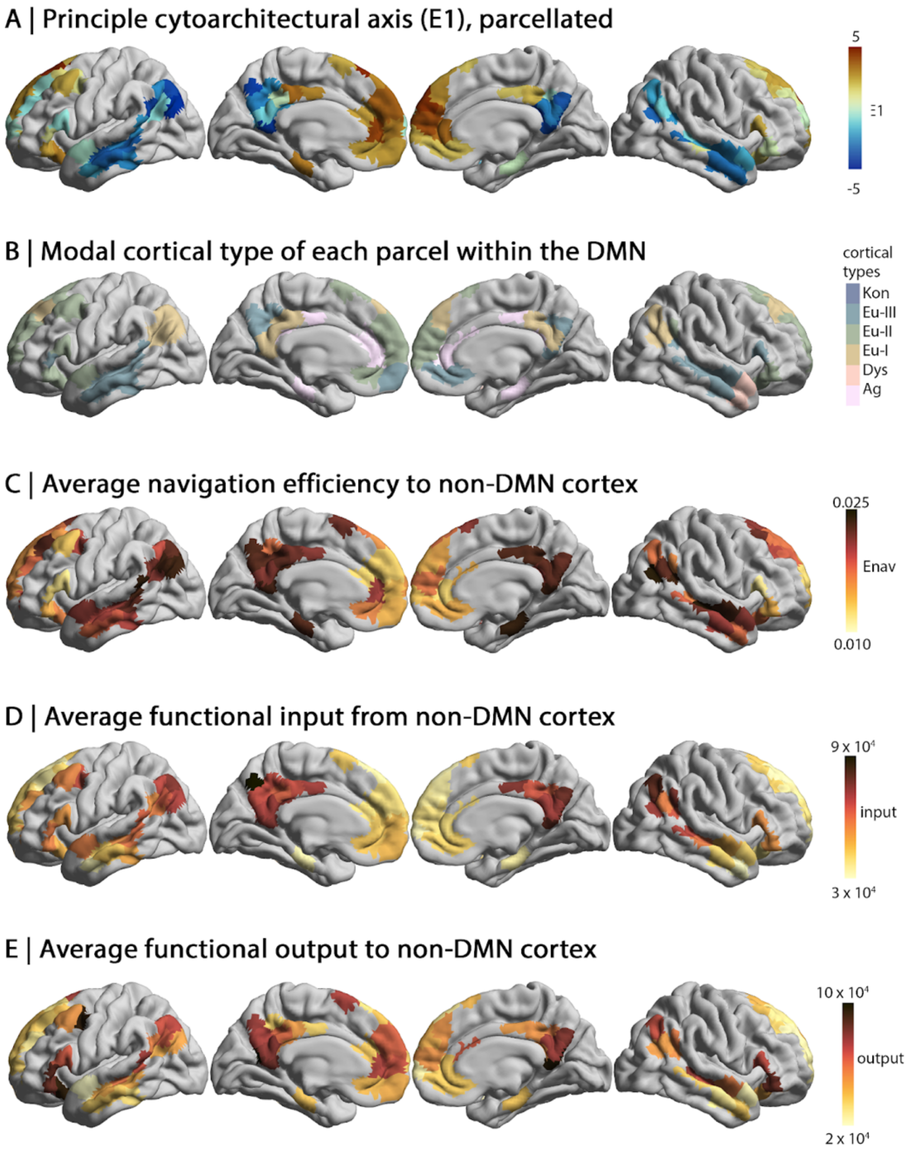


**Supplementary Figure 7:** Cortical maps illustrate the key axes of variation in A-B) cytoarchitecture, C) structural connectivity and D-E) signal flow. Exact values for each parcel can be found in Supplementary Table 1.

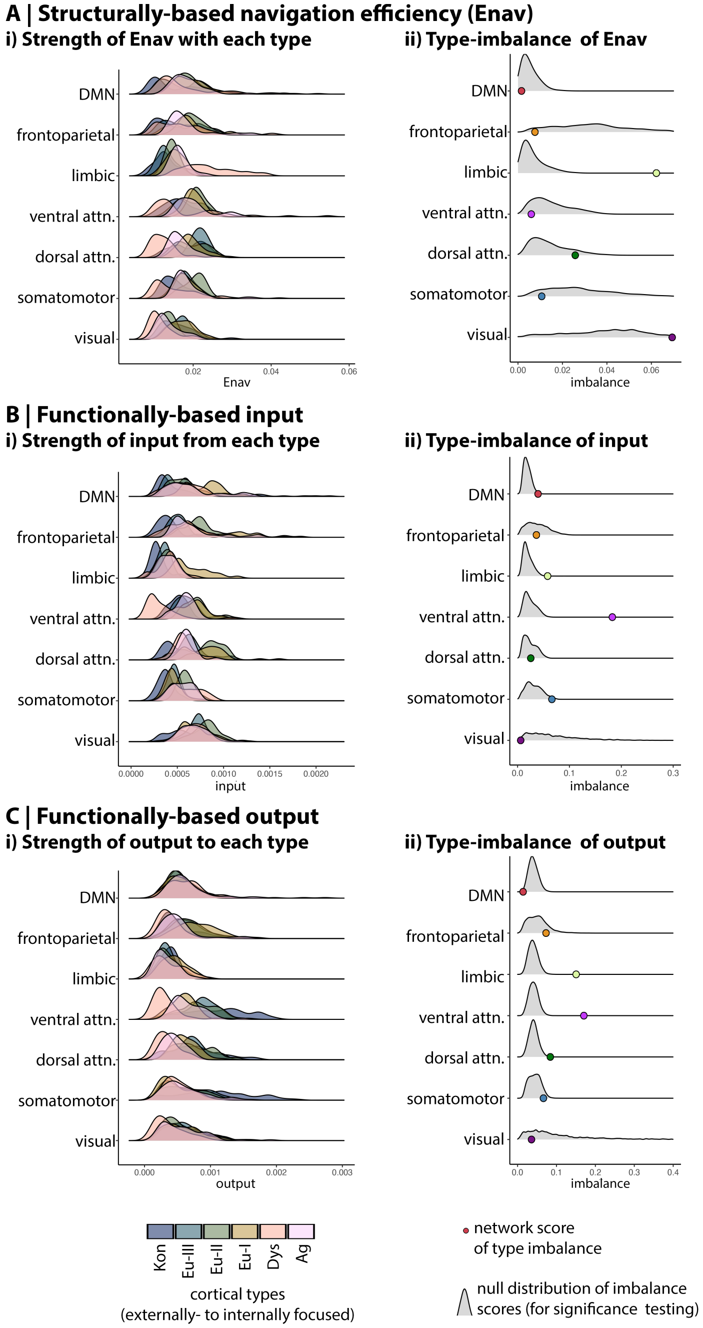


**Supplementary Figure 8: Comparison of functional networks based on inter-network connectivity to different cortical types.** Coloured ridge plots on the left of each panel show probability distributions of connectivity between the functional networks and non-DMN cortical types. We evaluated the imbalance of connectivity across cortical types using the Kullback-Leibler (KL) divergence from a null model with equal connectivity to each type. On the right of each panel, coloured dots show the empirical KL divergence for each network and the grey density plots show the null distribution of KL divergence values based on 10,000 spin permutations. **A)** The DMN exhibits the most balanced navigation efficiency across cortical types, compared to other functional networks. The balance of the DMN did not reach a level of significance relative to spin permutations, but spin permutations account for the size and distribution of the network, thus we may infer it is the large size and wide distribution of the network that enable the DMN to strike a balance in communication across cortical types. **B)** Input to the DMN is not balanced with regards to cortical types. Stronger input comes from heteromodal, Eu-I cortex, which aligns with the over-representation of this cortical type within the DMN. **C)** The DMN is unique amongst functional networks in exhibiting balanced output to all cortical types, which is further supported by the balance of the DMN reaching significance in spin permutation testing.

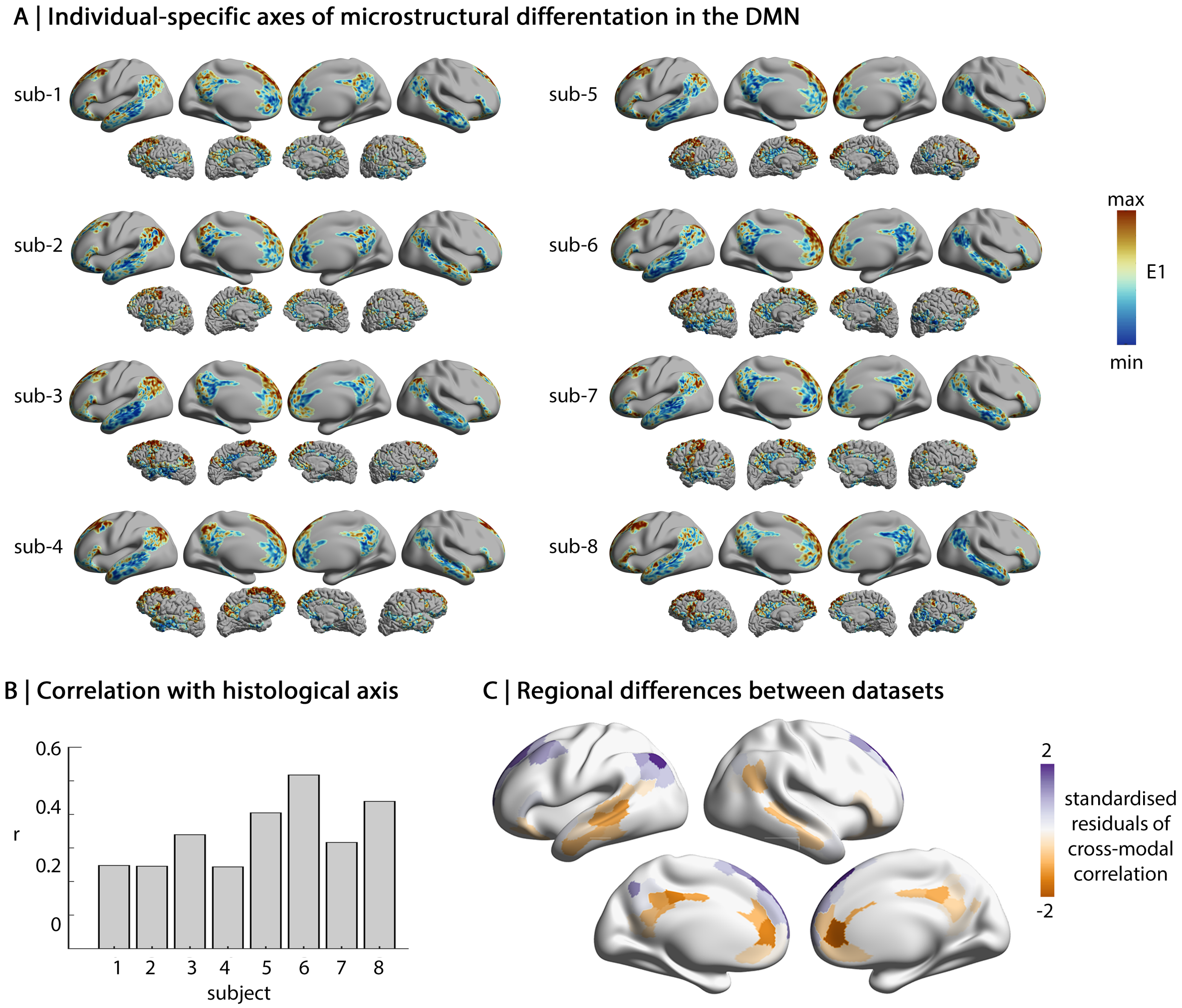


**Supplementary Figure 9: A)** The principal eigenvector of microstructural variation in the DMN (E1) extracted from myelin-sensitive quantitative MRI (qT1) is displayed for all eight subjects. For each subject, E1 is presented using atlas-based DMN on a standard surface (above) and individual-specific DMN on the subject's cortical surface (below). Notably, E1 was very similar between atlas- and individual-based reconstructions of the DMN. **B)** Subject-specific correlations of the qT1-derived axis with the axis of histological variation derived from BigBrain. **C)** Group-average standardised residuals highlight which parcels deviate from cross-modal alignment. Residuals were calculated by fitting a linear model between each subject-specific axis and the histological axis. Dark purple in the left lateral parietal area signifies higher E1 in the MRI dataset relative to the histological dataset, whereas dark orange in the anterior cingulate indicates lower E1 in the MRI dataset.

**Supplementary Table 1**

| E1 | Parcel name | Subregion | Von Economo | Cortical type | Enav | Afferent |
|---|---|---|---|---|---|---|
| -4.01 | 7Networks_LH_Default_Par_3 | Inferior parietal | PG | Eu-I | 2.0E-02 | 7.0E-04 |
| -3.80 | 7Networks_LH_Default_pCunPCC_1 | Precuneus | LC2 | Eu-I | 2.0E-02 | 7.3E-04 |
| -3.70 | 7Networks_LH_Default_pCunPCC_10 | Precuneus | PE | Eu-III | 2.0E-02 | 9.0E-04 |
| -3.63 | 7Networks_LH_Default_Par_5 | Inferior parietal | PG | Eu-I | 2.2E-02 | 5.6E-04 |
| -3.62 | 7Networks_RH_Default_pCunPCC_2 | Precuneus | LE | Agranular | 2.0E-02 | 6.9E-04 |
| -3.59 | 7Networks_RH_Default_pCunPCC_1 | Precuneus | OA | Eu-III | 2.1E-02 | 7.8E-04 |
| -3.53 | 7Networks_LH_Default_Par_6 | Inferior parietal | PG | Eu-I | 2.0E-02 | 7.5E-04 |
| -3.25 | 7Networks_RH_Default_pCunPCC_5 | Precuneus | PE | Eu-III | 2.0E-02 | 7.7E-04 |
| -3.14 | 7Networks_LH_Default_Temp_4 | Middle temporal | TE | Eu-III | 1.9E-02 | 4.6E-04 |
| -3.09 | 7Networks_LH_Default_Temp_6 | Middle temporal | TE | Eu-III | 2.0E-02 | 6.0E-04 |
| -3.06 | 7Networks_RH_Default_Par_2 | Inferior parietal | PG | Eu-I | 2.1E-02 | 7.3E-04 |
| -2.94 | 7Networks_LH_Default_Temp_2 | Middle temporal | TE | Eu-III | 1.7E-02 | 4.7E-04 |
| -2.89 | 7Networks_RH_Default_Temp_7 | Middle temporal | TE | Eu-III | 2.0E-02 | 6.9E-04 |
| -2.79 | 7Networks_RH_Default_Temp_2 | Middle temporal | TE | Eu-III | 1.6E-02 | 4.4E-04 |
| -2.74 | 7Networks_RH_Default_Temp_1 | Middle temporal | TG | Dysgranular | 1.5E-02 | 3.4E-04 |
| -2.58 | 7Networks_RH_Default_pCunPCC_4 | Precuneus | LC1 | Eu-I | 2.0E-02 | 7.1E-04 |
| -2.53 | 7Networks_LH_Default_pCunPCC_6 | Precuneus | PE | Eu-III | 2.0E-02 | 6.8E-04 |
| -2.48 | 7Networks_LH_Default_Temp_7 | Middle temporal | TE | Eu-III | 2.1E-02 | 5.6E-04 |
| -2.41 | 7Networks_LH_Default_Temp_3 | Middle temporal | TE | Eu-III | 1.8E-02 | 5.2E-04 |
| -2.40 | 7Networks_LH_Default_Par_7 | Inferior parietal | PG | Eu-I | 2.1E-02 | 6.8E-04 |
| -2.36 | 7Networks_RH_Default_Temp_4 | Middle temporal | TE | Eu-III | 1.9E-02 | 5.0E-04 |
| -2.31 | 7Networks_LH_Default_Temp_10 | Middle temporal | TE | Eu-III | 2.2E-02 | 6.2E-04 |
| -2.25 | 7Networks_LH_Default_pCunPCC_11 | Precuneus | PE | Eu-III | 2.1E-02 | 6.8E-04 |
| -2.15 | 7Networks_RH_Default_Par_1 | Inferior parietal | TE | Eu-III | 2.3E-02 | 6.6E-04 |
| -2.06 | 7Networks_LH_Default_Par_1 | Inferior parietal | TE | Eu-III | 2.2E-02 | 6.3E-04 |
| -1.91 | 7Networks_LH_Default_pCunPCC_4 | Precuneus | LE | Agranular | 2.0E-02 | 7.4E-04 |
| -1.84 | 7Networks_RH_Default_Par_5 | Inferior parietal | PG | Eu-I | 2.1E-02 | 7.9E-04 |
| -1.81 | 7Networks_LH_Default_pCunPCC_2 | Precuneus | PE | Eu-III | 2.2E-02 | 7.2E-04 |
| -1.70 | 7Networks_RH_Default_Temp_6 | Middle temporal | TE | Eu-III | 2.1E-02 | 6.2E-04 |
| -1.66 | 7Networks_LH_Default_Temp_1 | Middle temporal | TE | Eu-III | 1.6E-02 | 3.6E-04 |
| -1.58 | 7Networks_RH_Default_Temp_3 | Middle temporal | TG | Dysgranular | 1.7E-02 | 4.4E-04 |
| -1.05 | 7Networks_LH_Default_pCunPCC_3 | Precuneus | LC1 | Eu-I | 1.9E-02 | 7.1E-04 |
| -0.95 | 7Networks_RH_Default_Par_3 | Inferior parietal | PG | Eu-I | 2.2E-02 | 5.9E-04 |
| -0.95 | 7Networks_RH_Default_Temp_8 | Middle temporal | TE | Eu-III | 2.2E-02 | 6.9E-04 |
| -0.94 | 7Networks_LH_Default_Par_4 | Inferior parietal | PG | Eu-I | 2.2E-02 | 6.5E-04 |
| -0.76 | 7Networks_LH_Default_Par_2 | Inferior parietal | PG | Eu-I | 2.2E-02 | 5.9E-04 |
| -0.74 | 7Networks_LH_Default_PFC_15 | Superior frontal and ACC | FD | Eu-II | 1.5E-02 | 5.4E-04 |
| -0.64 | 7Networks_LH_Default_PFC_10 | Inferior frontal | FCBm | Eu-II | 1.9E-02 | 5.4E-04 |
| -0.53 | 7Networks_LH_Default_PFC_19 | Superior frontal and ACC | FC | Eu-I | 1.5E-02 | 3.8E-04 |
| -0.52 | 7Networks_LH_Default_PFC_16 | Superior frontal and ACC | FC | Eu-I | 1.7E-02 | 4.8E-04 |
| -0.46 | 7Networks_RH_Default_pCunPCC_8 | Precuneus | PE | Eu-III | 2.2E-02 | 6.4E-04 |
| -0.37 | 7Networks_LH_Default_PFC_11 | Superior frontal and ACC | FD | Eu-II | 1.2E-02 | 4.5E-04 |
| -0.30 | 7Networks_LH_Default_Temp_5 | Middle temporal | TA | Eu-II | 1.9E-02 | 5.9E-04 |
| -0.08 | 7Networks_RH_Default_Par_4 | Inferior parietal | PF | Eu-II | 2.3E-02 | 7.6E-04 |
| -0.05 | 7Networks_RH_Default_PFCdPFCm_6 | Superior frontal and ACC | LA2 | Agranular | 1.6E-02 | 4.3E-04 |
| 0.08 | 7Networks_RH_Default_PFCdPFCm_5 | Superior frontal and ACC | FD | Eu-II | 1.2E-02 | 4.2E-04 |
| 0.11 | 7Networks_RH_Limbic_TempPole_7 | Parahippocampal | HC | Agranular | 1.7E-02 | 3.8E-04 |
| 0.11 | 7Networks_RH_Default_pCunPCC_7 | Precuneus | LC1 | Eu-I | 2.1E-02 | 6.4E-04 |
| 0.28 | 7Networks_RH_Default_PFCv_3 | Inferior frontal | FF | Eu-II | 1.6E-02 | 5.5E-04 |
| 0.52 | 7Networks_RH_Default_PFCdPFCm_8 | Superior frontal and ACC | FD | Eu-II | 1.4E-02 | 3.8E-04 |
| 0.67 | 7Networks_RH_Default_Temp_5 | Middle temporal | TE | Eu-III | 1.9E-02 | 5.9E-04 |
| 0.69 | 7Networks_LH_Default_PFC_7 | Inferior frontal | FDT | Eu-III | 1.8E-02 | 6.0E-04 |
| 0.73 | 7Networks_LH_Default_PFC_20 | Superior frontal and ACC | FB | Eu-II | 2.1E-02 | 6.6E-04 |
| 0.77 | 7Networks_LH_Default_PFC_21 | Superior frontal and ACC | FB | Eu-II | 1.8E-02 | 5.3E-04 |
| 0.83 | 7Networks_RH_Default_PFCdPFCm_12 | Superior frontal and ACC | FB | Eu-II | 1.8E-02 | 4.5E-04 |
| 0.94 | 7Networks_LH_Default_PFC_22 | Superior frontal and ACC | FB | Eu-II | 1.8E-02 | 4.4E-04 |
| 0.96 | 7Networks_RH_Default_PFCv_1 | Inferior frontal | FF | Eu-II | 1.7E-02 | 4.0E-04 |
| 1.13 | 7Networks_LH_Default_PFC_6 | Superior frontal and ACC | FD | Eu-II | 1.4E-02 | 5.5E-04 |
| 1.16 | 7Networks_RH_Default_pCunPCC_7 | Precuneus | LA1 | Agranular | 2.1E-02 | 7.4E-04 |
| 1.18 | 7Networks_RH_Default_PFCdPFCm_13 | Superior frontal and ACC | FB | Eu-II | 1.6E-02 | 4.6E-04 |
| 1.27 | 7Networks_RH_Default_PFCdPFCm_1 | Superior frontal and ACC | FE | Eu-III | 1.4E-02 | 4.5E-04 |
| 1.31 | 7Networks_RH_Default_PFCdPFCm_10 | Superior frontal and ACC | FC | Eu-I | 1.7E-02 | 4.5E-04 |
| 1.40 | 7Networks_RH_Default_pCunPCC_6 | Precuneus | LC1 | Eu-I | 2.1E-02 | 6.5E-04 |
| 1.48 | 7Networks_LH_Default_PFC_17 | Superior frontal and ACC | FB | Eu-II | 1.8E-02 | 5.7E-04 |
| 1.48 | 7Networks_RH_Default_PFCv_4 | Inferior frontal | FDT | Eu-III | 1.9E-02 | 5.6E-04 |
| 1.49 | 7Networks_RH_Default_PFCdPFCm_2 | Superior frontal and ACC | FD | Eu-II | 1.1E-02 | 3.6E-04 |
| 1.50 | 7Networks_LH_Default_PFC_24 | Superior frontal and ACC | FB | Eu-II | 1.6E-02 | 4.6E-04 |

| | | | | | | |
|---|---|---|---|---|---|---|
| 1.64 | 7Networks_LH_Default_PFC_4 | Superior frontal and ACC | FH | Eu-II | 1.4E-02 | 4.7E-04 |
| 1.68 | 7Networks_RH_Default_PFCdPFCm_11 | Superior frontal and ACC | FC | Eu-I | 1.4E-02 | 3.6E-04 |
| 1.69 | 7Networks_LH_Default_PFC_12 | Superior frontal and ACC | LA1 | Agranular | 1.5E-02 | 4.3E-04 |
| 1.73 | 7Networks_LH_Default_PFC_5 | Inferior frontal | FF | Eu-II | 1.6E-02 | 5.8E-04 |
| 1.77 | 7Networks_LH_Default_PFC_3 | Superior frontal and ACC | FE | Eu-III | 1.2E-02 | 4.9E-04 |
| 1.82 | 7Networks_LH_Default_PFC_2 | Inferior frontal | FF | Eu-II | 1.6E-02 | 5.0E-04 |
| 1.83 | 7Networks_LH_Default_PFC_14 | Superior frontal and ACC | FD | Eu-II | 1.3E-02 | 4.4E-04 |
| 1.91 | 7Networks_RH_Default_PFCdPFCm_9 | Superior frontal and ACC | FC | Eu-I | 1.4E-02 | 3.8E-04 |
| 2.03 | 7Networks_LH_Default_pCunPCC_9 | Precuneus | LA1 | Agranular | 2.0E-02 | 7.1E-04 |
| 2.05 | 7Networks_RH_Default_PFCv_2 | Inferior frontal | FF | Eu-II | 1.6E-02 | 4.3E-04 |
| 2.09 | 7Networks_LH_Default_PFC_18 | Superior frontal and ACC | FC | Eu-I | 1.3E-02 | 3.8E-04 |
| 2.18 | 7Networks_LH_Default_PFC_1 | Inferior frontal | FF | Eu-II | 1.8E-02 | 5.0E-04 |
| 2.24 | 7Networks_LH_Default_PFC_8 | Superior frontal and ACC | FD | Eu-II | 1.2E-02 | 4.9E-04 |
| 2.29 | 7Networks_LH_Limbic_TempPole_8 | Parahippocampal | HC | Agranular | 1.7E-02 | 3.7E-04 |
| 2.58 | 7Networks_LH_Default_pCunPCC_8 | Precuneus | LC2 | Eu-I | 2.1E-02 | 6.7E-04 |
| 2.62 | 7Networks_RH_Default_PFCdPFCm_3 | Superior frontal and ACC | LA1 | Agranular | 1.4E-02 | 4.3E-04 |
| 2.77 | 7Networks_LH_Default_PFC_13 | Superior frontal and ACC | FD | Eu-II | 1.2E-02 | 4.2E-04 |
| 3.15 | 7Networks_RH_Default_PFCdPFCm_4 | Superior frontal and ACC | FD | Eu-II | 1.3E-02 | 4.2E-04 |
| 3.29 | 7Networks_LH_Default_PFC_9 | Superior frontal and ACC | LA1 | Agranular | 1.3E-02 | 4.6E-04 |
| 3.53 | 7Networks_RH_Default_PFCdPFCm_7 | Superior frontal and ACC | FD | Eu-II | 1.2E-02 | 3.9E-04 |
| 3.68 | 7Networks_LH_Default_PFC_23 | Superior frontal and ACC | FB | Eu-II | 1.5E-02 | 3.7E-04 |

**Supplementary Table 2: Cortical types by functional network**

| | Kon | Eu-III | Eu-II | Eu-I | Dys | Ag | Total vertices | KS statistic[1] |
|---|---|---|---|---|---|---|---|---|
| Visual | 0.29 | 0.41 | 0.17 | 0.10 | 0 | 0.03 | 2750 | 0.36, p<0.001 |
| Somatomotor | 0.10 | 0.54 | 0.31 | 0.04 | <0.01 | 0.01 | 3751 | 0.20, p<0.001 |
| DAN | <0.01 | 0.40 | 0.53 | 0.06 | 0 | 0 | 2188 | 0.29, p<0.001 |
| VAN | 0.02 | 0.18 | 0.50 | 0.13 | 0.08 | 0.09 | 2285 | 0.13, p<0.001 |
| Limbic | 0 | 0.24 | 0.28 | 0.11 | 0.26 | 0.10 | 1426 | 0.27, p<0.001 |
| Frontoparietal | 0 | 0.18 | 0.56 | 0.23 | <0.01 | 0.04 | 2314 | 0.11, p<0.001 |
| Default mode | <0.01 | 0.32 | 0.31 | 0.28 | 0.02 | 0.07 | 3765 | |
| Total vertices | 1218 | 6400 | 6805 | 2572 | 648 | 836 | | |

[1]Kolmogorov-Smirnov tests for independence of samples were calculated between each network and the DMN.

*Note:* entries in the centre of the table are proportions, which are provided relative to the functional network (ie: 29% of the visual network is koniocortical), thereby the rows approximately sum to 1 (given rounding errors).

Kon=koniocortical. Eu=eulaminate. Dys=dysgranular. Ag=agranular. DAN=dorsal attention network. VAN=ventral attention network.

**Supplementary Table 3: Correlation of DMN connectivity with cytoarchitectural axis**

| Measure of connectivity | Dataset | All non-DMN | Koniocortical | Eulaminate-III | Eulaminate-II | Eulaminate-I | Dysgranular | Agranular |
|---|---|---|---|---|---|---|---|---|
| $E_{NAV}$ (Structural model) | MICS | **r=-0.60, p<0.001** | **r=-0.63, p<0.001** | **r=-0.60, p<0.001** | r=-0.38, p=0.006 | r=-0.26, p=0.094 | r=0.09, p=0.291 | r=0.26, p=0.051 |
| | HCP | **r=-0.59, p<0.001** | **r=-0.43, p<0.001** | **r=-0.64, p<0.001** | r=-0.24, p=0.134 | r=-0.14, p=0.232 | r=0.08, p=0.328 | r=0.34, p=0.038 |
| Input (Functional model) | MICS | **r=-0.54, p<0.001** | **r=-0.38, p=0.006** | **r=-0.62, p<0.001** | **r=-0.35, p=0.005** | **r=-0.48, p=0.001** | r=-0.30, p=0.014 | **r=-0.43, p=0.003** |
| | HCP | **r=-0.40, p<0.001** | r=-0.23, p=0.085 | **r=-0.49, p<0.001** | **r=-0.33, p=0.003** | r=-0.20, p=0.044 | r=-0.21, p=0.015 | r=-0.18, p=0.140 |
| Output (Functional model) | MICS | r=-0.18, p=0.064 | r=-0.10, p=0.201 | r=-0.23, p=0.015 | r=-0.13, p=0.015 | r=-0.07, p=0.281 | r=-0.15, p=0.069 | r=-0.04, p=0.382 |
| | HCP | **r=-0.36, p<0.001** | **r=-0.30, p<0.001** | **r=-0.40, p=0.001** | r=-0.20, p=0.035 | r=-0.26, p=0.007 | r=-0.15, p=0.051 | r=-0.28, p=0.017 |
| Input (Extended functional model) | MICS | **r=-0.45, p<0.001** | **r=-0.42, p<0.001** | **r=-0.54, p<0.001** | **r=-0.22, p<0.001** | **r=-0.41, p<0.001** | r=-0.23, p=0.086 | r=-0.23, p=0.061 |
| | HCP | **r=-0.39, p<0.001** | **r=-0.28, p=0.004** | **r=-0.47, p<0.001** | r=-0.29, p=0.011 | r=-0.20, p=0.007 | r=-0.12, p=0.310 | r=-0.13, p=0.180 |
| Output (Extended functional model) | MICS | r=-0.12, p=0.200 | r=-0.18, p=0.131 | r=-0.02, p=0.035 | r=0.02, p=0.220 | r=-0.23, p=0.725 | r=0.04, p=0.058 | r=0.15, p=0.857 |
| | HCP | **r=-0.36, p<0.001** | **r=-0.32, p=0.001** | **r=-0.43, p<0.001** | r=-0.23, p=0.033 | **r=-0.21, p=0.004** | r=-0.12, p=0.240 | r=-0.19, p=0.061 |

*Note*: p-values reflect a two-sided comparison with 10,000 permutations. Significance (in bold) was deemed where p<0.004, which reflects a Bonferroni correction for seven two-side tests (each row of the table), with an alpha level of 0.05.

**Supplementary Table 4: Imbalance of connectivity across cortical types**

| Measure of connectivity | Dataset | Visual | Somato-motor | Dorsal attention | Ventral attention | Limbic | Fronto-parietal | Default mode |
|---|---|---|---|---|---|---|---|---|
| $E_{NAV}$ (Structural model) | MICS | KL=0.069, p=0.994 | KL=0.011, p=0.464 | KL=0.026, p=0.870 | KL=0.006, p=0.548 | KL=0.064, p=0.913 | KL=0.007, p=0.749 | KL=0.002, p=0.214 |
| | HCP | KL=0.089, p=0.741 | KL=0.022, p=0.282 | KL=0.057, p=0.752 | KL=0.016, p=0.352 | KL=0.083, p=0.520 | KL=0.014, p=0.442 | KL=0.004, p=0.104 |
| Input (Functional model) | MICS | KL=0.003, p=0.033 | KL=0.024, p=0.660 | KL=0.021, p=0.828 | KL=0.032, p>0.999 | KL=0.048, p>0.999 | KL=0.025, p=0.477 | KL=0.048, p=0.910 |
| | HCP | KL=0.033, p=0.411 | KL=0.032, p=0.809 | KL=0.017, p=0.548 | KL=0.092, p>0.999 | KL=0.019, p=0.677 | KL=0.039, p=0.822 | KL=0.022, p=0.827 |
| Output (Functional model) | MICS | KL=0.012, p=0.224 | KL=0.062, p=0.987 | KL=0.050, p>0.999 | KL=0.106, p>0.999 | KL=0.014, p=0.108 | KL=0.040, p=0.761 | **KL=0.003, p=0.001** |
| | HCP | KL=0.043, p=0.326 | KL=0.131, p>0.999 | KL=0.056, p=0.861 | KL=0.096, p>0.999 | KL=0.018, p=0.128 | KL=0.031, p=0.423 | **KL=0.004, p<0.001** |
| Input (Extended functional model) | MICS | KL=0.013, p=0.221 | KL=0.017, p=0.695 | KL=0.004, p=0.999 | KL=0.064, p>0.999 | KL=0.040, p=0.924 | KL=0.029, p>0.999 | KL=0.013, p=0.841 |
| | HCP | KL=0.111, p=0.869 | KL=0.092, p=0.695 | KL=0.116, p=0.978 | KL=0.194, p>0.999 | KL=0.036, p=0.052 | KL=0.091, p=0.834 | KL=0.045, p=0.001 |
| Output (Extended functional model) | MICS | KL=0.040, p=0.513 | KL=0.051, p=0.887 | KL=0.085, p>0.999 | KL=0.108, p>0.999 | KL=0.022, p=0.209 | KL=0.061, p>0.999 | **KL=0.008, p<0.001** |
| | HCP | KL=0.056, p=0.337 | KL=0.158, p>0.999 | KL=0.117, p=0.978 | KL=0.150, p>0.999 | KL=0.029, p=0.078 | KL=0.073, p=0.612 | **KL=0.032, p<0.001** |

*Note:* p-values reflect a one-sided comparison with 10,000 permutations. Significance (in bold) was deemed where p<0.007, which reflects a Bonferroni correction for seven one-side tests (tests within a row of the table), with an alpha level of 0.05.
